# *Drosophila* midgut tumor-induced insulin resistance systemically remodels lymph gland hematopoiesis during cancer cachexia

**DOI:** 10.1101/2025.11.24.690188

**Authors:** Ujjayita Chowdhury, Gauri Panzade, Priya Neelesh Karnik, Pushkar Birwadkar, Tashu Tashu, Rohan Jayant Khadilkar

**Affiliations:** Stem cell and Tissue Homeostasis lab, Cancer Research Institute, ACTREC – Tata Memorial Centre, Navi Mumbai – 410210, India; Homi Bhabha National Institute, Anushaktinagar, Mumbai – 400094, India

**Keywords:** Cancer cachexia, insulin resistance, signaling, hematopoiesis, *Drosophila*

## Abstract

Cancer cachexia involves systemic metabolic deregulation along with classical features of muscle wasting, lipolysis, and chronic inflammation. While tumor non-autonomous effects on peripheral organs are recognized, how the tumor rewires the circulating immune cells and hematopoiesis remains unclear. We utilized a *Drosophila* larval cancer cachexia model by expressing *yki^3SA^* in the adult midgut precursors (AMP), which gives rise to a tumor in the larval midgut and recapitulates key cachectic phenotypes, including insulin resistance. Tumor-induced cachexia results in perturbed blood cell homeostasis with a reduced niche and aberrant blood cell differentiation in the larval hematopoietic organ, the lymph gland (LG). Bulk RNA-seq analysis of circulating hemocytes from tumor-bearing larvae revealed upregulation of multiple cachectic ligands, notably ImpL2, an insulin antagonist. We demonstrate that elevated ImpL2 levels reduce LG niche size and promote aberrant blood cell differentiation. Elevated ImpL2 levels and systemic insulin resistance in the tumor-induced cachexia conditions result in abrogation of insulin signaling in the niche-progenitor micro-environment in the LG. DE-cadherin levels in the primary LG lobe are perturbed, and Wingless signaling is down-regulated, driving prohemocyte differentiation. A genetic mimic of systemic ImpL2 overexpression or high sugar diet (HSD) conditions recapitulates these LG phenotypes due to abrogation of the Insulin-Wingless signaling axis. Hemocyte-specific ImpL2 depletion in HSD-fed larvae rescued these defects, suggesting a regulatory role for circulating hemocyte-derived ImpL2. Our findings reveal that hemocyte-derived factors actively contribute to systemic insulin resistance, causing hematopoietic remodeling in cancer cachexia.

## Introduction

Tumors actively communicate with their surroundings through intricate molecular signaling pathways involving neighboring cells and the extracellular matrix^1–3^. Tumor expansion and metastatic potential are profoundly impacted by the local micro-environment that delivers essential nutrients to the tumor^2,3^. While the role of tumor micro-environment as a key driver of cancer progression has been well explored, emerging evidence reveals an equally important but less appreciated paradigm: the tumor macro-environment^4,5^. This refers to the far-reaching, systemic crosstalk between malignant cells and distant organs throughout the body. There is a molecular dialogue between the tumor and peripheral host organs, in which the tumor hijacks normal physiological processes in remote tissues, leading to devastating consequences^5,6^. When this inter-organ communication becomes severely disrupted, patients may develop complex metabolic disorders, most notably cancer-associated cachexia^7–10^. This debilitating syndrome manifests as progressive muscle wasting coupled with dysregulated energy expenditure - symptoms that persist despite adequate nutritional intervention. Cachexia affects approximately 80% of patients with solid tumors and directly contributes to 20-30% of cancer-related deaths annually^10–14^.

Systemic inflammation is central to the pathophysiology of cancer cachexia. Tumors release pro-inflammatory cytokines, including Tumor Necrosis Factor-alpha (TNF-α), Interleukin-6 (IL-6), and Interleukin-1-beta (IL-1β), among many other molecules that circulate systemically^8–10^. These factors reach distant organs such as the adipose tissue, skeletal muscle, bone, liver, and brain, where they disrupt normal metabolic function, compromise tissue homeostasis, and trigger widespread metabolic abnormalities^10^. Additionally, this inflammatory state impairs insulin signaling in peripheral tissues, establishing tumor-induced insulin resistance^15–17^. This metabolic reprogramming represents a maladaptive response that simultaneously compromises host physiology while creating conditions favorable for tumor progression.

The influence of immune cells on cancer progression has been extensively documented ^18–22^. Hematopoiesis—the physiological process of blood cell formation—depends on carefully orchestrated extrinsic and intrinsic signals that maintain stem cell pools and guide differentiation into mature blood lineages with immune function^23,24^ . Under normal conditions, these regulatory mechanisms work together to preserve stem cell and tissue homeostasis. Few pieces of evidence indicate that epithelial tumors can significantly perturb hematopoietic function^25–27^. Hematological abnormalities have emerged as important systemic consequences of cancer, yet they remain inadequately understood^28^. Tumor-secreted factors can rewire bone marrow dynamics and stimulate acute myelopoiesis, suggesting these hematological changes may extend beyond mere secondary effects of malignancy^29^. The developmental mechanisms underlying these alterations, however, warrant further investigation to fully elucidate their contribution to cancer pathophysiology. *Drosophila* is a well-established system for modelling various kinds of cancers and to study tumor-host interactions^30–34^. To mechanistically dissect how epithelial tumors systemically influence hematopoiesis at a distant site, we employed *Drosophila* which has a dedicated hematopoietic organ in the larval stage.

In *Drosophila* larvae, blood cell formation occurs primarily in the primary lobe of the lymph gland (LG), which is subdivided into distinct regions: the posterior signaling center (PSC), a hematopoietic stem cell niche that serves as a regulatory hub; the medullary zone (MZ), which contains undifferentiated blood cell precursors known as prohemocytes; and cortical zone (CZ) which houses mature blood cells^35–38^ . Recent single-cell sequencing of the LG hemocytes has comprehensively revealed the extensive heterogeneity of hemocyte subtypes within the LG^39–41^. Advanced technologies have elucidated the previously under-characterized roles of diverse prohemocyte populations, including their functions in lipid metabolism, stress responses, and development^39^. Flies harbor three distinct hemocyte types: plasmatocytes, the most abundant type that performs phagocytic functions by engulfing pathogens similar to vertebrate macrophages; crystal cells participate in wound healing and melanization immune responses; and lamellocytes are specialized cells that combat parasitic wasp infestation^35–38^. *Drosophila* hematopoiesis has been extensively studied under various developmental and disease conditions, revealing that prohemocytes can sense nutritional signals and modulate multiple signaling pathways to maintain the balance between self-renewal and differentiation into mature lineages^38,42–44^

Innate immune responses in flies can be classified into two major categories: the humoral immune response by innate immune organs like the fat body, gut, trachea, and the hemocytes which produce anti-microbial peptides, while the cellular immune response is largely mediated by hemocytes^45–49^. Circulating hemocytes perform diverse functions, including extracellular matrix secretion, phagocytosis of pathogens, and antimicrobial peptide production^50–53^. Emerging evidence also demonstrates a functional role for hemocytes in tumor responses^37,54–59^.Given that circulating hemocytes constitute a key component of the systemic environment, it would be interesting to understand whether epithelial tumors can rewire or educate these circulating cells and whether that would benefit the tumor or not. These circulating hemocytes could play an important regulatory role in modulating the extent of cachexia, resource mobilization to and from the tumor to distant tissues, and control the innate immune status. An open, unexplored question is whether these circulating hemocytes or factors produced by them can remodel resident hematopoiesis in the LG in such conditions.

Here, we demonstrate that the aberrant proliferation of adult midgut precursors (AMPs) driven by *yki^3SA^* expression in the larval midgut induces cachexia-like symptoms and peripheral insulin resistance. Using this larval tumor-induced cachectic model, we observe that LG niche size reduces with an aberrant increase in blood cell differentiation in the cachexia background. We also find that elevated ImpL2 levels from both the larval gut tumor and uniquely even from circulating hemocytes cause insulin resistance systemically. Our results indicate that insulin signaling in the niche is perturbed, leading to a decrease in PSC/niche size. Reduced niche size and abrogated insulin signaling impacts Wingless signaling activity in the progenitors, thereby affecting DE-cadherin levels in the progenitors, pushing them towards differentiation. Our observations indicate that elevated ImpL2 levels cause insulin resistance systemically, which perturbs the niche-progenitor micro-environment, impacting blood cell homeostasis in the larval LG. Our study provides important insights into the inter-organ crosstalk between the tumor, circulating blood cells, and the LG in cachexia conditions. The increased blood cell differentiation is very similar to skewed myelopoiesis in vertebrates in a tumor background, which could give rise to chronic systemic inflammation, a characteristic feature of cancer cachexia that aggravates the syndrome. This study reveals mechanistic insights into how cellular immune response is heightened in cancer cachexia, with important clinical implications. Exploring ways of reversing the onset of systemic inflammation has significant promise in ameliorating cancer cachexia, which needs further investigation.

## Results

### *yki^3SA^* - induced larval gut tumor causes cachectic phenotypes

A constitutively active form of Yorkie, (*yki^3SA^*, *yki^S111A-S168A-S250A^* triple mutant), when expressed in the intestinal stem cells (ISCs) in the adult midgut, is known to cause over-proliferation of ISCs^31,60^,ultimately causing tumors and peripheral organ wasting^31,61,62^. To induce larval midgut tumors, we expressed *yki^3SA^* in the adult midgut precursors (AMPs) using *esg-Gal4, UAS-GFP*. *esg-Gal4* mediated expression of *yki^3SA^* resulted in an expansion of the Esg-GFP cluster volume, i.e., the Esg-GFP positive AMP cell population, as compared to the wild type control (Fig. 1A-B). These *yki^3SA^-*driven AMP tumors in the larval midgut resulted in the breakdown of Coracle-positive septate junctions, which appeared discontinuous and disrupted as compared to the wild type control (Fig. 1C-D), establishing that the *yki^3SA^* mutation in the larval midgut potently drives tumors.

**Fig 1:**
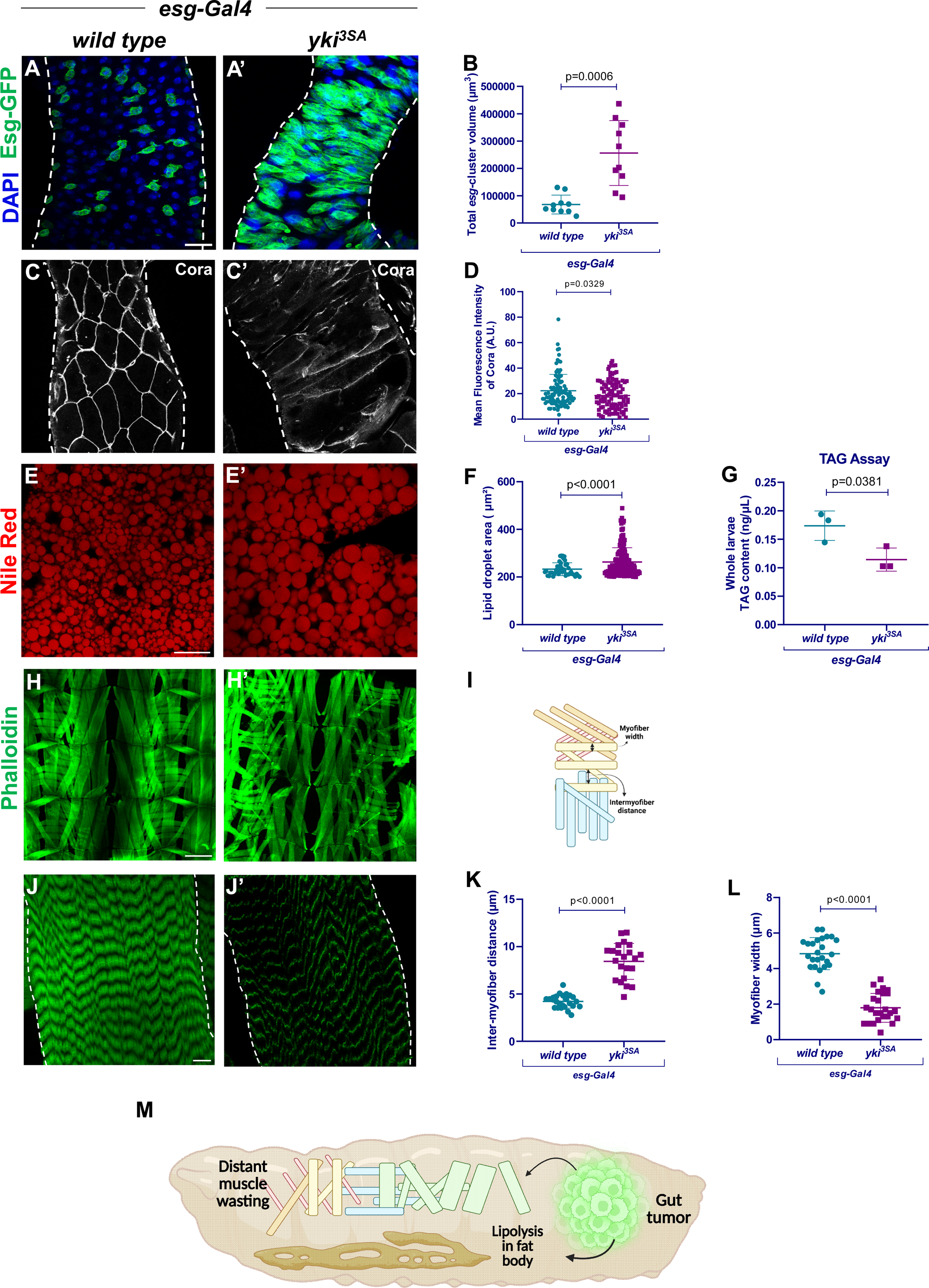
*yki^3SA^*-induced larval midgut tumor causes cachectic phenotypes. Adult midgut progenitor (AMP)- specific expression of the oncogenic mutation, *yki^3SA^* in the larval midgut using *esg-Gal4, UAS-GFP,* compared to the wild type control (A-A’). Quantitation of the GFP-positive Esg-cluster volume in larvae bearing *yki^3SA^*-induced tumor as compared to control (B). Larval midguts stained with Coracle (Gray), a septate junction marker in larvae bearing *yki^3SA^*- induced tumor as compared to the wild type control (C-C’). Corresponding graph of quantitation of the Mean Fluorescence Intensity of Coracle (D). Larval fat body lipid droplets (LDs) stained with Nile Red (Red) in larvae bearing *yki^3SA^*-induced tumors as compared to the wild type control (E-E’). Estimation of the corresponding lipid droplet area of LDs from larval fat bodies (F). Quantification of whole larval Triacylglycerol (TAG) content in larvae bearing tumor (*yki^3SA^*) compared to the control (G). Larval cuticular muscle stained with Phalloidin (Green) in larvae bearing *yki^3SA^*-induced tumor compared to the wild type control (H-H’; J-J’). Schematic illustration of parameters for the quantitation of the width of the muscle fillet (I). Created in BioRender. Khadilkar, R. (2025) https://BioRender.com/3655czf. Quantitation of myofiber width (K) and inter-myofiber distance (L) of the muscle fillet of larvae bearing *yki^3SA^*-induced tumor compared to the wild type control. Schematic illustration of larval gut tumor showing distant wasting effects (M). Created in BioRender. Khadilkar, R. (2025) https://BioRender.com/e36k015. *esg-Gal4, UAS-GFP* crossed to *Canton-S* was used as the wild type. Nuclei were stained with DAPI (Blue). White dotted lines indicate the border of the larval midgut and muscle fillet, respectively. Scale Bar: 50μm (A-A’, C-C’), 40μm (E-E’), 200μm (H-H’), 10μm (J-J’). Statistical analysis was performed using Student’s t-test with Welch’s correction. Values are displayed as Mean ± SD. p-values indicating statistical significance have been mentioned in the respective graphs.

We then investigated the fat bodies of the larvae bearing *yki^3SA^*-induced midgut tumors, and stained them with a lipophilic dye, Nile Red, which showed an accumulation of lipid droplets (LDs) as compared to the wild type control (Fig. 1E-E’). Quantitation of LD area revealed that the LD area was overall higher in the fat bodies from the larvae bearing the midgut tumors (Fig. 1F). This phenotype is also observed in other cachexia models reported earlier, which suggests mobilization of resources and an efflux of lipids into the hemolymph^32,63–65^. Additionally, the larvae bearing the *yki^3SA^* -*induced* midgut tumors display visual bloating, a striking hallmark of the tumor-induced wasting phenotype (Fig. S1A). To characterize LD size distribution, we generated a heat-map displaying LD area across larval fat bodies. This visualization revealed that tumor-bearing larvae possessed enlarged LDs relative to wild type larvae (Fig. S1B). To assess the whole-body Triacylglycerol (TAG) content, we measured the total TAG content of the whole larvae and found that there is a significant decrease in the TAG content in the larvae bearing *yki^3SA^*-induced tumors (Fig. 1G). We then tested the other major hallmark of cachexia, i.e., muscle wasting, wherein we visualized the larval cuticular muscle fibers using Phalloidin and observed that there is a muscle thinning phenotype. It was observed that the thickness of each myofiber in the larvae bearing the midgut tumor is less than the wild type larvae, whereas the distance between each myofiber increases, showing a muscle thinning phenotype (Fig. 1H-L). Our observations indicate that *yki^3SA^* induction in the larval midgut also drives characteristic tumor-induced wasting phenotypes in host organs like muscles and adipose tissue that are major hallmarks of cancer-induced cachexia (Fig. 1M).

### *yki^3SA^* - induced larval gut tumors result in systemic insulin resistance

Tumors can alter systemic metabolism through secreted factors^4,66,67^. To investigate potential metabolic perturbations in the host tissue, we assessed insulin signaling, a critical regulator of glucose uptake and metabolism, in the larval fat body. To test this, we checked if *yki^3SA^* midgut tumors systemically alter insulin sensitivity in peripheral tissues like the fat body, which is a major nutrient-sensing hub^68^. We observed that the tumor-bearing larvae exhibit a striking decrease in pAkt levels, which is a readout of insulin signaling in the fat bodies as compared to control larvae (Fig. 2A-B’, E). We then performed the 2-NBDG (a glucose analog) uptake assay in the larval fat bodies of the *yki^3SA^* tumor-bearing larvae and observed a decreased uptake of 2-NBDG as compared to the control, implying reduced glucose uptake in the fat body (Fig. 2C-D’, F). Taken together, these results suggest that there is a severe disruption of insulin signaling in the fat body in the larval cachexia model.

**Fig 2:**
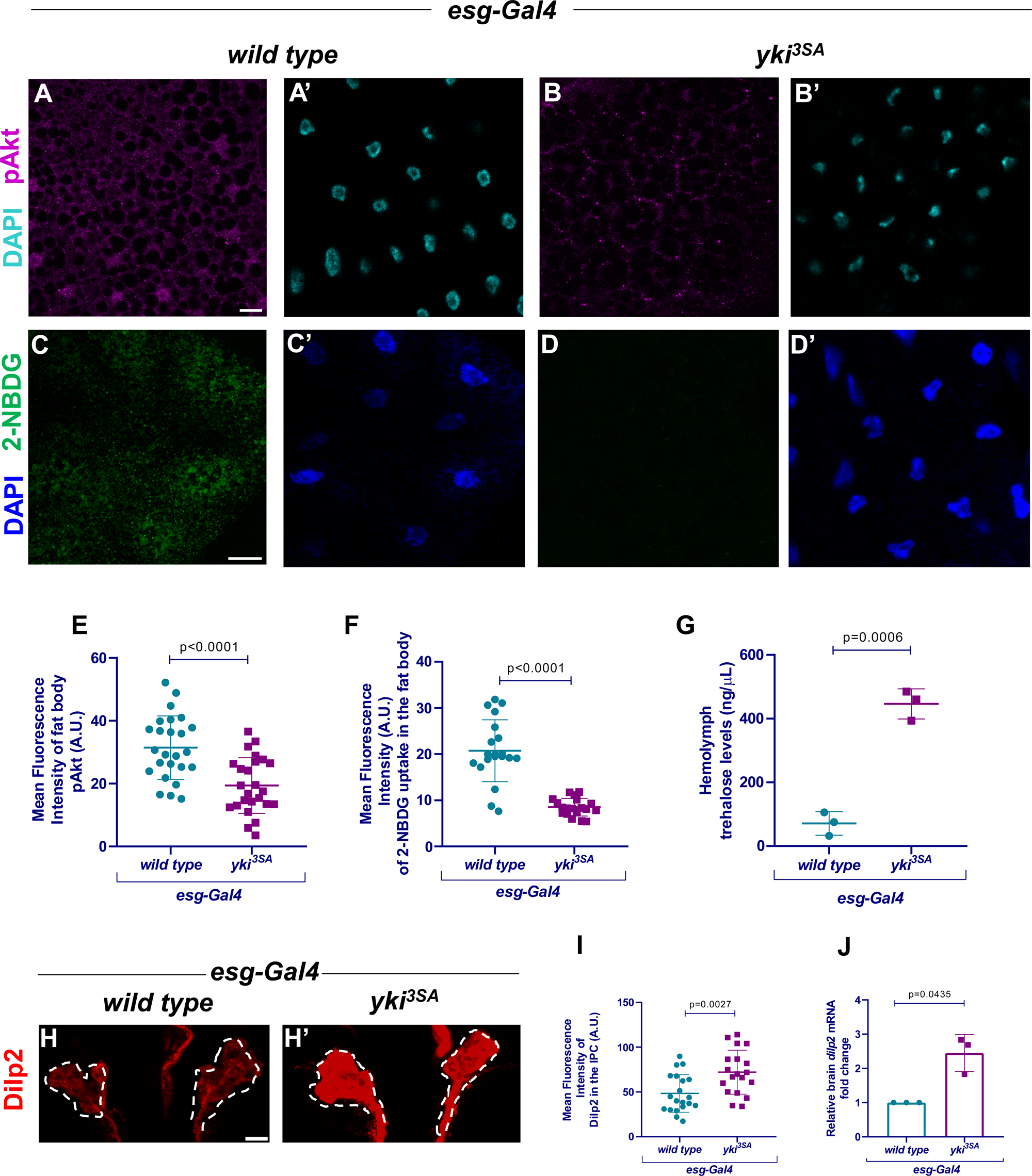
*yki^3SA^*-larval midgut tumor results in systemic insulin resistance. Larval fat body stained with pAkt (Magenta) from larvae bearing tumor (*yki^3SA^*) compared to the wild type control (A-B’). 2-NBDG (Green) uptake in the fat body in larvae bearing a tumor (*yki^3SA^*), compared to wild type control (C-D’). Graphs showing quantitation of the Mean Fluorescence Intensity of pAkt (E) or 2-NBDG (F) in the fat body of larvae bearing tumor (*yki^3SA^*) compared to wild type control. Circulating trehalose levels in the hemolymph of the larvae bearing tumor (*yki^3SA^*) compared to the wild type control (G). Larval insulin-producing cells (IPCs) stained with Dilp2 (Red) of larvae-bearing tumor (*yki^3SA^*) compared to the wild type control (H-H’). Corresponding graph showing quantitation of the Mean Fluorescence Intensity of Dilp2 levels (I) and relative *dilp2* mRNA expression (J) in the larval IPCs. *esg-Gal4, UAS-GFP* crossed to *Canton-S* was used as the wild type. White dotted lines indicate the border of the IPCs in the larval brain. Nuclei were stained with DAPI (Cyan in A’, B’; Blue; in C’, D’). Scale Bar: 20μm (A-D’), 10μm (H-H’). Statistical analysis was performed using Student’s t-test with Welch’s correction. Values are displayed as Mean ± SD. p-values indicating statistical significance have been mentioned in the respective graphs.

Given how closely cachexia and insulin resistance are tied^7,15–17,69,70^, we hypothesized that the larval gut tumor results in an insulin resistance-like condition. Hence, we checked circulating trehalose levels in the larval hemolymph and observed a marked increase in trehalose levels in the cachectic larvae, implicating hyperglycemia typical of insulin resistance (Fig. 2G). Quite expectedly, immunostaining of the larval insulin-producing cells (IPCs) in the brain with Dilp2 (*Drosophila* insulin-like peptide 2) showed higher Dilp2 intensity in the cachectic larvae as compared to the wild type larvae and higher *dilp2* mRNA levels in the larval brain of the cachectic larvae as compared to wild type (Fig. 2H-J). Altogether, these results demonstrate reduced insulin sensitivity, hyperglycemia, and hyperinsulinemia, all of which contribute to insulin resistance in the larval cachexia model.

### LG blood cell homeostasis is perturbed in larvae bearing *yki^3SA^* -induced midgut tumors

Chronic systemic inflammation is a hallmark of cancer cachexia^8,10^. Hematopoiesis, the process of formation of blood or immune cells, is tightly linked to inflammation^71–74^ . Epithelial tumors have been reported to disrupt hematopoiesis by remodeling the bone marrow niche and causing myeloid skewing^27,28^. Despite growing evidence linking tumors to systemic inflammation, the mechanisms by which tumor-derived signals disrupt hematopoiesis and contribute to chronic inflammatory states remain incompletely characterized. Here, we wanted to understand how the *yki^3SA^* induced midgut tumors impact hematopoiesis in the *Drosophila* larval hematopoietic organ, the lymph gland (LG). Besides, several reports have shown that the LG is a sensitive organ that responds to systemic signals like starvation and stress^37,38,42,75–77^ . We studied the LGs from tumor-bearing larvae for various hematopoietic parameters. We checked the LG whole mounts and found that there is no disruption of LGs observed in tumor-bearing larvae as compared to the wild type control (Fig. S2A-A’), and corresponding quantitation indicated no significant change in the area of the primary lobe of LGs (Fig. S2B). A close examination of the PSC region marked by Antennapedia (Antp) revealed that third instar tumor-bearing larvae exhibit a marked decrease in Antp-positive niche cells as compared to the wild type third instar larvae (Fig. 3A-B). We then visualized the medullary zone progenitors using DE-cadherin and found that the typical DE-cadherin cortical pattern observed in wild type larval LGs is disrupted and lost in the LGs from tumor-bearing larvae, indicating loss of progenitors in the LGs from tumor-bearing larvae (Fig. 3C-D).

**Fig 3:**
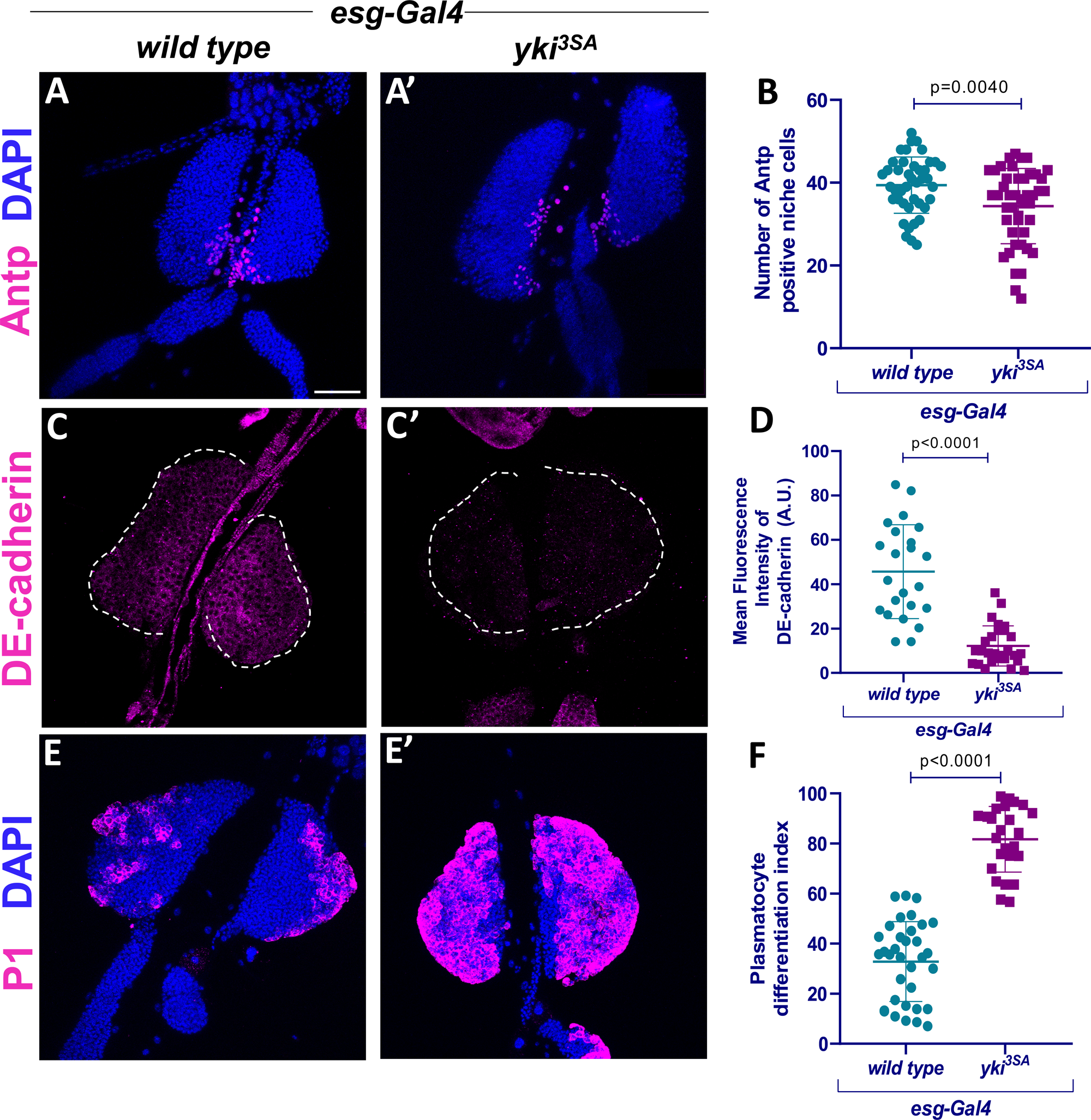
*yki^3SA^* larval midgut tumor exerts systemic effects that disrupt lymph gland hematopoiesis. Larval primary lymph gland (LG) lobes stained with Antp (Magenta; A-A’), DE-cadherin (Magenta; C-C’), or P1 (Magenta; E-E’) in larvae bearing *yki^3SA^* induced midgut tumors as compared to the wild type control. Quantitation of Antp-positive PSC/niche cells (B), Mean fluorescence intensity denoting DE-cadherin levels in the primary LG lobes (D), and plasmatocyte differentiation index in the primary LG lobes (F). *esg-Gal4, UAS-GFP* crossed to *Canton-S* was used as the wild type. Nuclei were stained with DAPI (Blue). White dotted lines indicate the border of the primary LG lobe. Scale Bar: 50μm (A-A’, C-C’, E-E’). Statistical analysis was performed using Student’s t-test with Welch’s correction. Values are displayed as Mean ± SD. p-values indicating statistical significance have been mentioned in the respective graphs.

The reduction in niche cells and the progenitor pool corroborated with the profile of blood cell differentiation, wherein we observed that there is an increase in NimRodC1 (P1) positive plasmatocyte differentiation (Fig. 3E-F), Hindsight positive crystal cells (Fig. S2C-E), and there is presence of ꞵ-integrin positive lamellocytes (Fig. S2F-H) in the LGs in larvae bearing the *yki^3SA^* induced midgut tumors as compared to the control. Together, these results demonstrate that LG hematopoiesis is indeed remodeled and perturbed in tumor-induced cachexia conditions, as there is a surge in terminally differentiated cells in the LG, which could be an indication of a heightened cellular immune response to the presence of the tumor.

### Circulating hemocytes from tumor-bearing larvae show altered gene signatures

Based on the LG phenotypes, we were intrigued to understand if tumor-secreted or other systemic factors impact LG blood cell homeostasis. Multiple studies in both larval and adult models of cachexia have profiled ligands, prominently ImpL2, Upd3, Bnl, and Pvf1 ^31,32,61,62,65,78,79^, which are upregulated in the tumors in a cachexia scenario in both larval and adult models. However, it is unclear whether circulating hemocytes in the larval hemolymph are also rewired or educated by the tumor and if their gene signatures are altered in a cancer cachexia scenario. We wanted to assess the extent to which the gene signatures in the circulating hemocytes were altered. Hence, we performed a bulk RNA-seq of the larval hemocytes in cachectic and wild type larvae (Fig. 4A). Differential gene expression analysis of the data shows 1009 differentially upregulated genes and 686 downregulated genes (Fig. S3A-C). PathON^62^, a bioinformatic tool that maps various signaling pathways and their ligands in bioinformatics data sets, helped us mine specific ligands upregulated in the hemocytes in our gut tumor-induced larval cachexia model (Fig. 4B, S3D). We observe few signaling ligands like Spitz, Spaetzle, Pyramus, Pvf2 etc. to be upregulated. Notably, we observed that ImpL2, a cachectic ligand, shows an average of 1.7-fold increase in larval hemocytes. Analysis of the differentially expressed gene sets showed a list of biological processes, cellular components, and molecular functions enriched, and dot plot visualization of Gene Set Enrichment Analysis suggested activated and suppressed pathways in the RNA-seq data (Fig. S3E-F). We focused on investigating ImpL2 further given the insulin resistance conditions and its unexplored role in regulating hematopoiesis. We validated the transcriptomics data using qPCR (Fig. 4C). We then wanted to see if there is any upregulation in the tumorous guts as well, and quite expectedly, we saw, the larval gut has a higher gene expression of ImpL2 as well (Fig. 4D). These data imply that ImpL2 is not only upregulated in the tumor alone but peripheral cells like circulating hemocytes also show an upregulation of ImpL2.

**Fig 4:**
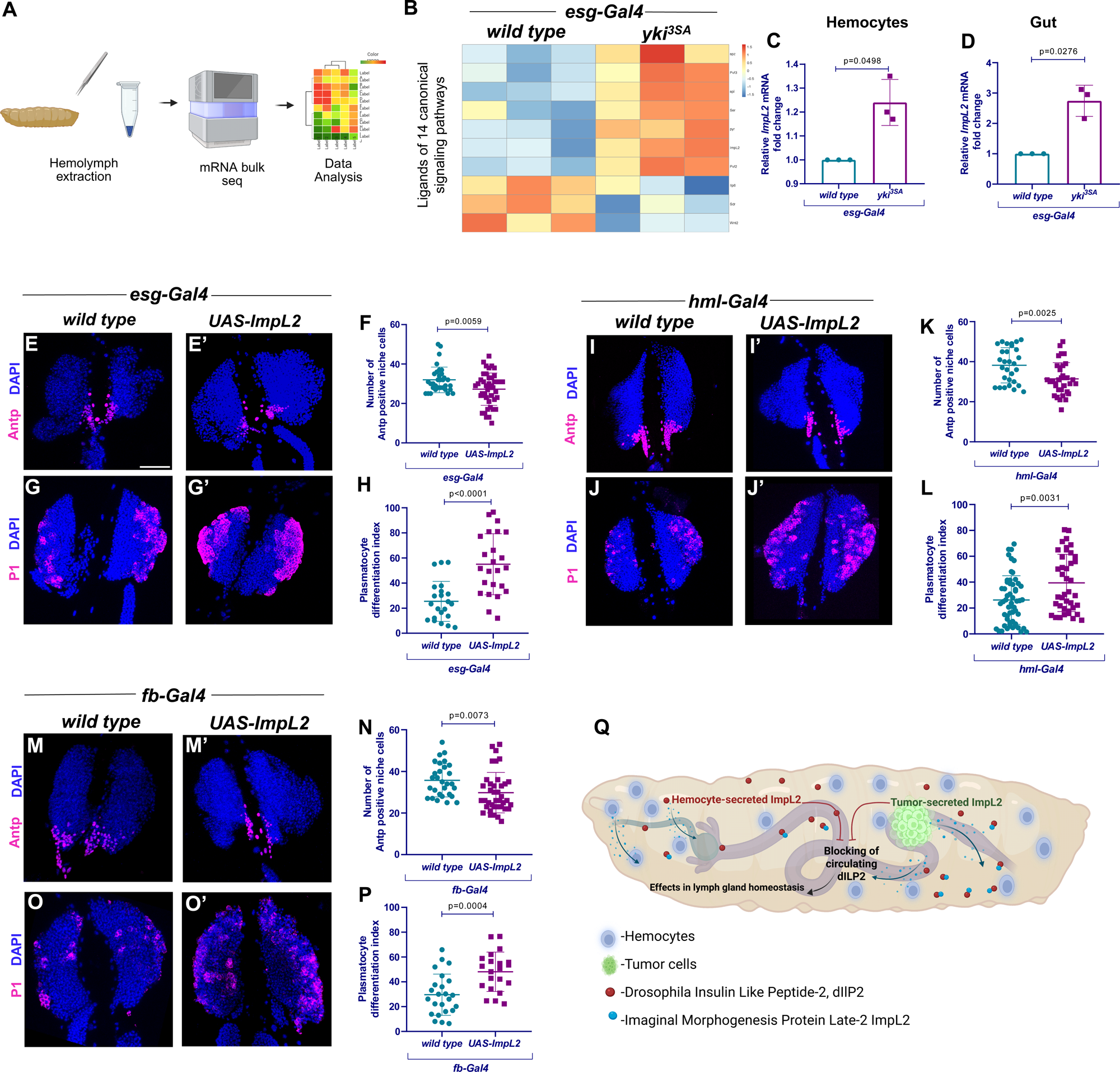
Systemic elevation of ImpL2 levels perturbs LG blood cell homeostasis. Schematic illustration of hemocyte isolation and bulk RNA sequencing workflow (A). Created in BioRender. Khadilkar, R. (2025) https://BioRender.com/l33r955. Heatmap visualization of ligands (of 14 canonical signaling pathways as listed in PathON) differentially expressed in transcriptomics of circulating hemocytes from larvae bearing tumor (*yki^3SA^*) compared to the wild type control (B). Relative *ImpL2* mRNA expression in the hemocytes and gut from larvae bearing tumor (*yki^3SA^*) compared to wild type control, respectively (C-D). Antp (Magenta) positive niche cell numbers or P1 (Magenta) positive plasmatocyte differentiation upon overexpression of ImpL2 using *esg-Gal4* in larval midgut AMPs (E’, G’) or using *hml-Gal4* in hemocyte population (I’, J’) or *fb-Gal4* in the fat body (M’, O’) as compared to respective controls- *esg-Gal4 X wt* (E, G) or *hml-Gal4 X wt* (I, J) or *fb-Gal4 X wt* (M, O). Quantitation of Antp-positive niche cell numbers in LG lobes upon overexpression of ImpL2 using the above Gal4s as compared to the control (F, K, and N). Plasmatocyte differentiation index upon ImpL2 overexpression using the above Gal4s as compared to the control (H, L, and P). Schematic showing the model of systemic ImpL2-driven effects in disrupting LG blood cell homeostasis in the larva bearing midgut tumor (Q). Created in BioRender. Khadilkar, R. (2025) https://BioRender.com/s53w321. Nuclei were stained with DAPI (Blue). Scale Bar: 50μm (E-G’, I-J’, M-O’). Statistical analysis was performed using Student’s t-test with Welch’s correction. Values are displayed as Mean ± SD. For transcriptomic analysis, hemocytes from 100 larvae were used for each biological replicate. Samples were prepared in triplicate. p-values indicating statistical significance have been mentioned in the respective graphs.

### An excess of systemic ImpL2 impairs LG blood cell homeostasis

Next, we wanted to investigate whether excess ImpL2 alone can cause a similar disruption in blood cell homeostasis as in the cancer cachexia condition. Hence, we designed genetic mimic experiments that can recapitulate the excess ImpL2 by overexpressing ImpL2 in various systemic organs like the gut, fat body, or circulating hemocytes alone and observed that there is a decrease in the number of Antp positive niche cells and an increase in plasmatocyte differentiation as compared to their respective controls (Fig. 4E-P), indicating that systemic upregulation of ImpL2 is critical in regulating resident blood cell homeostasis in the LG. Imaginal Morphogenesis Protein Late 2 (ImpL2) is classically known to bind Dilp2, thereby antagonising the insulin signaling pathway by making Dilp2 unavailable for Insulin receptor (InR) binding and signaling ^79^ (Fig. 4Q). Interestingly, we also observed that both *dilp2* mutant alleles *dilp2,3* and *dilp2,3,5* phenocopy LG phenotypes observed upon systemic ImpL2 overexpression. LGs of *dilp2,3* and *dilp2,3,5* mutants stained with Antp show a reduction in Antp-positive niche cell numbers (Fig. S4A-D) and an aberrant increase in plasmatocyte differentiation index (Fig. S4E-H). In line with these results, it was observed that IPC-specific overexpression of ImpL2 (*dilp2>UAS-ImpL2*) or knockdown of Dilp2 (*dilp2>UAS-dilp2RNAi*) resulted in a marked increase in plasmatocyte differentiation, corroborating the data for systemic ImpL2 overexpression (Fig. S4I-L).

### ImpL2 is capable of regulating hematopoiesis in a localized manner

Single-cell transcriptomic analysis demonstrates that ImpL2 is expressed in the LG as well as in distinct circulating hemocyte populations^39,41,50^. Also, an increase in systemic levels of ImpL2 triggered a plasmatocyte differentiation response; hence, we wanted to ascertain the localized effect of ImpL2 overexpression in the blood progenitor population. Core progenitor-specific overexpression of ImpL2 in the larval LGs (*tepIV>UAS-ImpL2*) exhibited an increased plasmatocyte differentiation index (Fig. S5A-C) as compared to the control. We also performed a pan-progenitor Gal4-specific overexpression of ImpL2 in the larval LGs (*domeMESO>UAS-ImpL2*) and observed an aberrant increase in plasmatocyte differentiation index recapitulating the systemically elevated ImpL2 phenotype (Fig. S5D-F). Since data mined from the Fly Hemocyte Atlas suggests that the PSC is enriched for ImpL2 transcripts, we performed a knockdown of ImpL2 in the niche region. Niche-specific knockdown of ImpL2 in the larval LGs (*antp-mRFP>ImpL2RNAi*) also showed increased plasmatocyte differentiation index (Fig. S5G-I), establishing the fact that even cell-autonomously ImpL2 is pertinent in regulating blood cell homeostasis in a localized manner.

### Tumor-derived systemic signals disrupt lymph gland homeostasis by driving aberrant differentiation through niche-progenitor insulin signaling impairment

Next, we wanted to probe the mechanism by which blood cell homeostasis is affected in the LGs of cachectic larvae. Strikingly, insulin signaling is essential for maintaining blood cell homeostasis in *Drosophila*, as it regulates both the hematopoietic niche and progenitor cells within the LG^42,43,77^. Both cell types respond to the insulin-like peptide Dilp2, which is critical for maintaining niche-progenitor equilibrium^38,42^ . Since one of the striking phenotypes for metabolic dysfunction in our larval cancer cachexia model is insulin resistance (Fig. 2), we asked whether insulin signaling in the LG from tumor-bearing larvae is affected or not. We first checked insulin sensitivity and activation status using a tGPH (a genetic PI3K reporter line) reporter^80,81^ . tGPH localizes to the membrane upon activation of the insulin signaling pathway and is cytoplasmic when inactive. We combined the tGPH reporter in the genetic background of the *yki^3SA^*-induced midgut tumor and found that there is higher cytoplasmic expression of tGPH in the Antp-positive niche cells of the LGs from the tumor-bearing larvae as compared to the control (Fig. 5A-C), indicating insulin pathway inactivation. Similarly, tGPH was quantified in the medullary zone progenitors (MZ) of the LGs, which showed that there is higher cytoplasmic expression of tGPH in the MZ progenitor region (Fig. 5D-F) flanking the dorsal vessel (DV), indicating low insulin signaling. Additionally, to validate the status of insulin signaling in the niche and MZ progenitors, we checked pAkt, a readout of active insulin signaling pathway. We co-stained the LGs with Antp and pAkt and observed that the levels of pAkt are lower in the Antp-positive niche cells, indicating low insulin signaling, thus validating the tGPH results in the tumor-bearing larvae as compared to the control (Fig. 5G-I). We also checked the status in the progenitors by co-staining with the plasmatocyte marker, P1, along with pAkt, and quantified the levels of pAkt in the non-P1 region that consists of non-differentiated blood cells or progenitors. We observed a decrease in pAkt levels in the progenitor region in LG from tumor-bearing larvae as compared to the control (Fig. 5J-L). Hematopoietic progenitors in the LG respond to systemic insulin signals and, in turn, activate Wnt/Wingless (Wg) signaling, which is essential for progenitor maintenance^42^. Also, Wingless signaling was reported to play a dual role in blood progenitor maintenance and crystal cell differentiation in the LG^82^. To further ascertain if Wg signaling is perturbed in the LGs in larvae with cancer induced cachexia; we stained the LGs for the Wg ligand and observed that there is a stark decrease in the expression of Wg in the hematopoietic progenitors of the *yki^3SA^* tumor-bearing larvae as compared to the control (Fig. 5M-O), indicating that the Wg signaling also goes down in the progenitors.

**Fig. 5:**
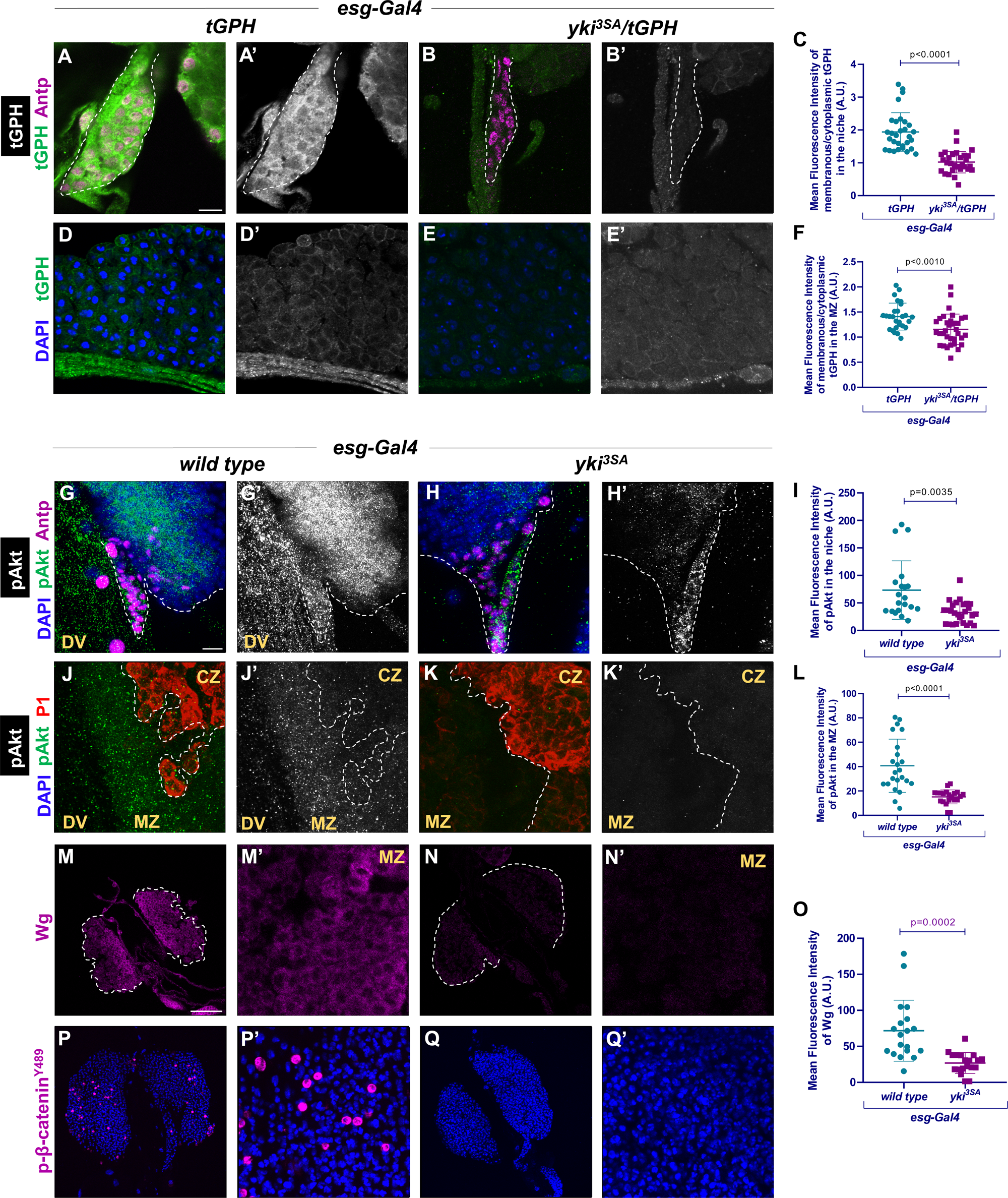
*yki^3SA^*-driven midgut tumor-derived systemic factors result in impairment of niche-progenitor insulin signaling. tGPH, an insulin signaling genetic reporter expression (Green; Grayscale) in the niche (A-B’) and MZ progenitors (D-E’) of the LG upon induction of *yki^3SA^* midgut tumors as compared to the wild type control. Quantitation of the Mean Fluorescence Intensity of the membranous: cytoplasmic ratio of tGPH in the niche (C) or the MZ (F) of the LG of larvae bearing tumor (*yki^3SA^*) compared to the wild type control. Primary LG lobe co-stained with pAkt (Green; Grayscale) and Antp (Magenta) marking the niche of larvae bearing tumor (*yki^3SA^*) compared to the wild type control (G-H’). Quantitation of the Mean Fluorescence Intensity of the pAkt levels in the niche of the LG of larvae bearing tumor (*yki^3SA^*) compared to the wild type control (I). Primary LG lobe co-stained with pAkt (Green; Grayscale) and P1 (Red); with P1-negative region denoting medullary zone (MZ) flanking the dorsal vessel (DV) of the LG of larvae bearing tumor (*yki^3SA^*) compared to the wild type control (J-K’). Corresponding graph showing quantitation of the mean fluorescence intensity of the pAkt levels in the MZ of the LG (L). Primary LG lobe stained with Wg (Magenta) (M-N’) or p-β-catenin^Y489^ (Magenta) (P-Q’) in the LG of larvae bearing tumor (*yki^3SA^*) compared to the wild type control. Quantitation of the mean fluorescence intensity of the Wg levels in the MZ in the LG of larvae bearing tumor (*yki^3SA^*) compared to the wild type control (O). *esg-Gal4, UAS-GFP* crossed to *+/+;tGPH/tGPH* was used as the control for (A-F). *esg-Gal4, UAS-GFP* crossed to *Canton-S* was used as the control for (G-O). Nuclei were stained with DAPI (Blue). Scale Bar: 10μm (A-B’, D-E’, G-H’, J-K’, M’, N’, P’, Q’), 50μm (M, N, P, Q). The white dotted line indicates the border of the niche (A-B’), (A-B’, G-H’), or demarcates differentiated plasmatocytes from the MZ (J-K’) or the border of the LG (M, N). Statistical analysis was performed using Student’s t-test with Welch’s correction. Values are displayed as Mean ± SD. p-values indicating statistical significance have been mentioned in the respective graphs.

We examined the readout wherein nuclear translocation of phosphorylated β-catenin (pY489-β-cat) serves as a hallmark of active Wnt/Wg signaling, occurring when the destruction complex is inhibited and stabilized β-catenin accumulates in the nucleus. This phenomenon is typically observed in hematopoietic progenitors, where a recent study^83^ demonstrated nuclear pY489-β-cat localization in the distal medullary zone and intermediate progenitors. To validate Wg signaling activation status, we immunostained samples with pY489-β-cat antibody and observed a striking reduction in nuclear β-catenin within hematopoietic progenitors of *yki^3SA^* larvae as compared to the control (Fig. 5P-Q’). Collectively, these findings demonstrate that tumor-derived systemic signals rewire hematopoietic homeostasis through a cascade of disruptions: inhibition of insulin signaling in both the niche and progenitors, suppression of the Wnt/Wg pathway in progenitors, compromising their maintenance, and promoting aberrant differentiation.

### ImpL2 depletion from the tumor restores lymph gland homeostasis in an insulin signaling-dependent manner

Since we observed that *yki^3SA^*-induced larval gut tumor shows upregulated ImpL2 levels, we were particularly intrigued to investigate if the tumor-secreted ImpL2 plays a role in the disruption of blood cell homeostasis in the LG in an insulin signaling-dependent manner. We hence depleted ImpL2 (*UAS-ImpL2RNAi*) in the *esg-Gal4* driven *yki^3SA^* gut tumor and examined the LGs compared to the parental control (*esg>ImpL2RNAi*) and the tumor control (*esg>yki^3SA^*). We stained the LGs with Antp and observed that there is a rescue in the Antp-positive niche cell numbers (Fig. 6A-D). We then checked the plasmatocytes and found a rescue in the aberrant plasmatocyte differentiation in the tumor-bearing larvae with ImpL2 knockdown in the tumor as compared to the tumor alone control (Fig. 6E-H). These LGs were stained with a progenitor marker, DE-cadherin, which also showed a rescue in DE-cadherin levels in the LG, indicating rescue of the progenitor pool upon depletion of ImpL2 from the tumor as compared to the tumor alone control (Fig. 6I-L). To investigate if the rescue is insulin signaling dependent, we co-stained the tumor-bearing larvae where ImpL2 is depleted from the tumor with Antp and pAkt, and observed that there is a rescue of pAkt levels in the niche co-stained with Antp as compared to the tumor alone control. (Fig. 6M-P). We were then curious to know if the insulin activity in MZ progenitors also shows a rescue, for which we also quantified pAkt levels in the MZ flanking the dorsal vessel and observed that there was a similar rescue of pAkt levels in the prohemocytes as well (Fig. 6Q-T). This implies that higher systemic ImpL2 is sufficient to affect and perturb insulin signaling in the hematopoietic compartments, like niche and MZ progenitors of the LG, thereby impacting blood cell homeostasis. Interestingly, when stained with Wg, we did not find a significant rescue in the Wg levels in the tumor-bearing larvae with ImpL2 knockdown in the tumor as compared to the tumor alone (Fig. 6U-X) implying that the rescue in insulin signaling is not enough to cause a rescue in the Wg signaling pathway and other alternate mechanisms might be playing a role in regulating the Wg pathway in the progenitors.

**Fig 6:**
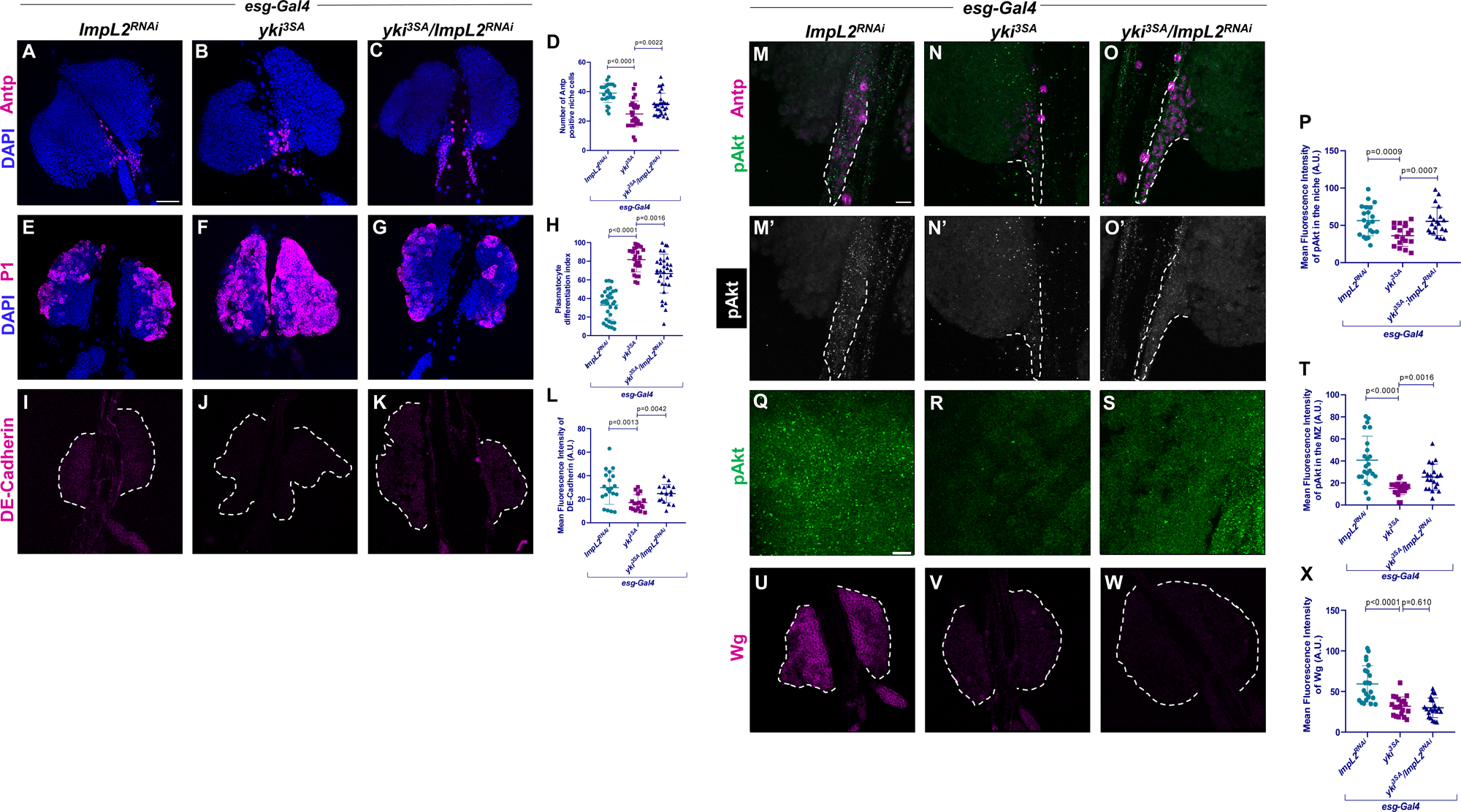
ImpL2 depletion from midgut tumor restores LG blood cell homeostasis via insulin signaling. Primary LG lobes stained with Antp (Magenta; A-C) or P1 (Magenta; E-G) or DE-cadherin (Magenta; I-K) upon depletion of ImpL2 in the genetic background of expression of *yki^3SA^* using *esg-Gal4, UAS-GFP* (C, G, K) as compared to the *yki^3SA^* induction using *esg-Gal4, UAS-GFP* tumor control (B, F, J) or the *ImpL2RNAi* driven by *esg-Gal4, UAS-GFP* control (A, E, I). Quantitation of Antp positive niche cell numbers (D), Plasmatocyte differentiation index (H), and mean fluorescence intensity of DE-cadherin expression in primary LG lobe (L). Primary LG lobe stained with pAkt (Green; Grayscale) in the Antp (Magenta) positive niche or in the MZ flanking the DV upon depletion of ImpL2 in the genetic background of expression of *yki^3SA^* using *esg-Gal4, UAS-GFP* (O, O’, S) as compared to the *yki^3SA^* mediated tumor induction using *esg-Gal4, UAS-GFP* (tumor control; N, N’, R) or the *ImpL2RNAi* driven by *esg-Gal4, UAS-GFP* control (M, M’, Q). Corresponding mean fluorescence intensity of pAkt in the niche (P), or MZ (T) in the primary LG lobe. Primary LG lobes stained with Wg (Magenta) upon depletion of ImpL2 in the genetic background of expression of *yki^3SA^* using *esg-al4, UAS-GFP* (W) as compared to the *yki^3SA^* induction using *esg-Gal4, UAS-GFP* tumor control (V) or the *ImpL2RNAi* driven by *esg-Gal4, UAS-GFP* alone control (U). Corresponding mean fluorescence intensity of the levels of Wg in the MZ of LG (X). Nuclei were stained with DAPI (Blue). Scale Bar: 50μm (A-C, E-G, I-K, U-W), 10μm (M-O’, Q-S). The white dotted line indicates the border of the LG (I-K, U-W), or the border of the niche (M-O’). Statistical analysis was performed using Student’s t-test with Welch’s correction. Values are displayed as Mean ± SD. p-values indicating statistical significance have been mentioned in the respective graphs.

In addition to investigating whether the LG phenotypes are rescued upon ImpL2 depletion in the tumor, we also assessed whether ImpL2 depletion has any effect on the tumor itself. Our observations indicate that there is no significant change in the Esg-GFP cluster volume in the midgut upon ImpL2 depletion as compared to the tumor alone (Fig. S6A-D). We next checked if the cachexia phenotypes are rescued, namely the lipid droplet area in the fat bodies and cuticular muscle parameters. We found that the fat body lipid droplet area shows a rescue along with an increase in the myofiber width of the cuticular muscles upon ImpL2 depletion in the *yki^3SA^* -induced midguts as compared to the tumor alone control (Fig. S6E-L).

### Hemocyte-mediated ImpL2 depletion rescues LG hematopoietic defects in a High Sugar Diet-induced insulin resistance model

Since transcriptomics analysis of the circulating hemocytes of *yki^3SA^* tumor-bearing larvae indicated an upregulation of ImpL2, we wanted to delineate whether hemocyte-derived ImpL2 affects LG blood cell homeostasis. We showed that the *yki^3SA^* tumor-bearing larvae display insulin resistance. Hence, we mimicked insulin resistance by feeding wild type larvae with HSD (High Sugar Diet, 1M sucrose). HSD-fed larvae are classically known to cause insulin resistance^84^. In our study as well, we validated the model by observing fat bodies of larvae fed with HSD stained with pAkt showed lower expression of pAkt, indicating low insulin sensitivity in the fat body as compared to the fat bodies of larvae fed with CD (Control Diet, 0.15M sucrose) (Fig. S7A-B, E). Additionally, the brains of larvae fed with HSD stained with Dilp2 antibody showed higher intensity in the HSD condition compared to CD (Fig. S7F-G, J), indicating hyperinsulinemia, which is typical of insulin resistance. Since we observed that tumor-bearing larvae had elevated ImpL2 levels, we wanted to investigate if HSD-induced insulin resistance causes systemic upregulation of ImpL2. Also, it is reported that ImpL2 is not produced in the larvae in standard conditions^85^; however, it was interesting to observe that HSD causes upregulation of ImpL2 not only systemically in whole larvae but also particularly in the circulating hemocytes (Fig. 7A-A’). However, depleting ImpL2 from the hemocytes (*hml>UAS-ImpL2RNAi*) in larvae fed with HSD is not able to rescue insulin sensitivity in the fat body, and the higher Dilp2 in the IPCs of the brain (Fig. S7C, E, H, J). Another striking observation is when we over-express ImpL2 in the hemocytes (*Hml>UAS-ImpL2*) and rear these flies in CD, we find that the pAkt levels in the fat bodies of such larvae are lower as compared to wild type fed with CD (Fig. S7D, E) and as in the case of the HSD insulin resistance phenotype there is higher Dilp2 expression in the larval brains in Hml-driven overexpression of ImpL2 fed with CD compared to wild type fed with CD (Fig. S7 I, J), implying ImpL2 is capable of driving systemic insulin resistance when over-expressed in the circulating hemocyte population.

**Fig 7:**
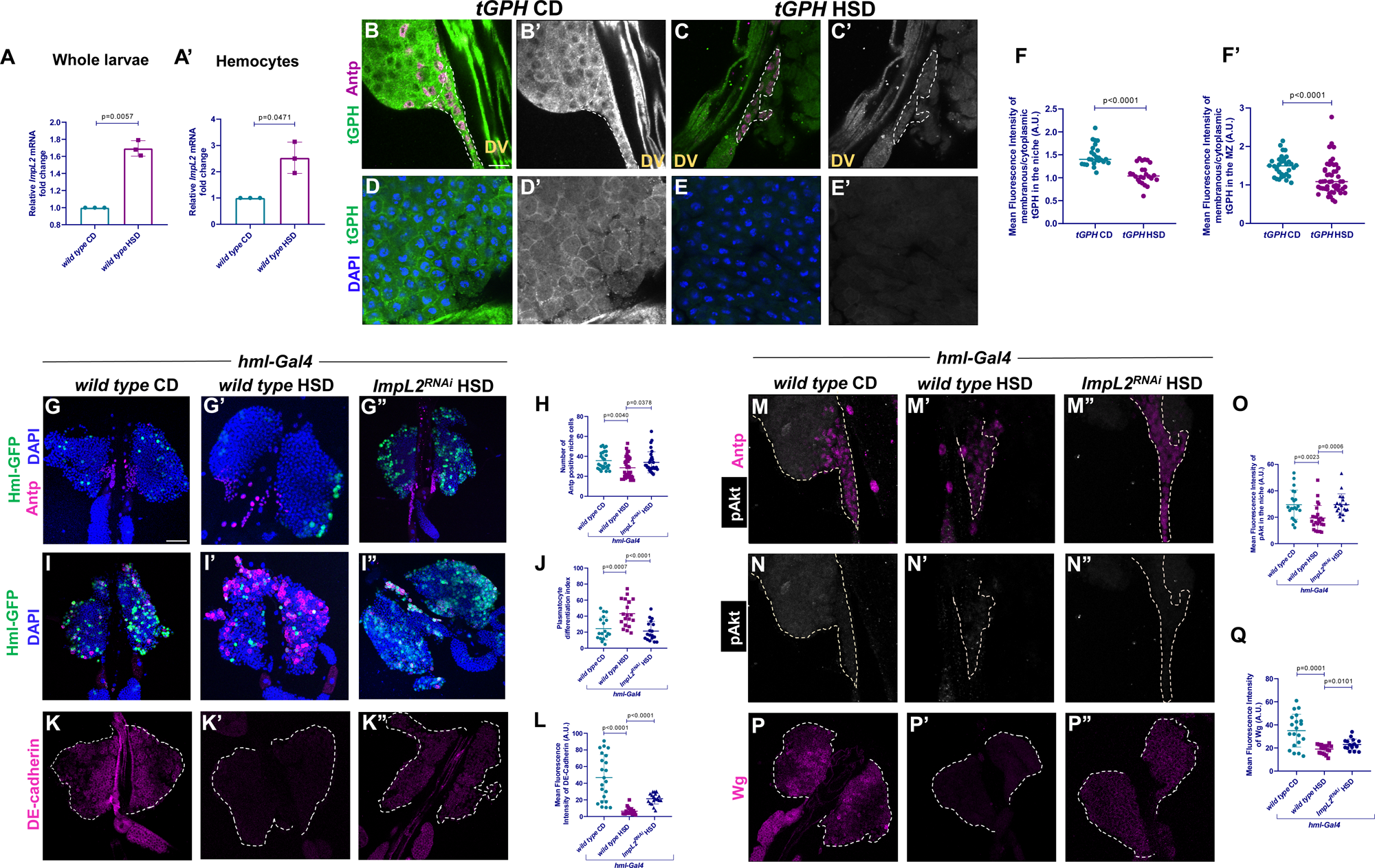
Hemocyte-specific ImpL2 depletion rescues LG hematopoiesis defects in a High Sugar Diet-induced insulin resistance model. Relative mRNA expression of *ImpL2* in whole larvae (A) or circulating hemocytes (A’) from High Sugar Diet (HSD) fed larvae compared to a Control Diet (CD) fed larvae. tGPH, an insulin signaling genetic reporter expression (Green; Grayscale) in the niche marked with Antp (Magenta) (B-C’) and MZ progenitor flanking the DV (D-E’) in the LG of the larvae fed with HSD compared to CD. Corresponding graph showing quantitation of the mean fluorescence intensity of the membranous to cytoplasmic intensity of tGPH in the niche and the medullary zone, respectively (F-F’). Larval LGs stained with Antp (Magenta) or P1 (Magenta) or DE-cadherin (Magenta) or pAkt (Grayscale) co-stained with Antp (Magenta) to mark the niche or Wg (Magenta) in the HSD-fed larvae with *hml-Gal4, UAS-GFP* (Green) driven knockdown of ImpL2 (G’’, I’’, K’’, M’’, N’’, P’’) compared to HSD-fed larvae (G’, I’, K’, M’, N’, P’) and CD-fed larvae (G, I, K, M, N, P). Corresponding graph showing the quantitation of the number of Antp positive niche cells (H), or Plasmatocyte Differentiation Index (J), or mean fluorescence intensity of DE-cadherin (L), or pAkt (O), or Wg (Q) in the respective genotypes/groups. *Canton-S* was used as the *wild type*. Nuclei were stained with DAPI (Blue). Scale Bar: 50μm (G-G’’, I-I’’, K-K’’, P-P’’), 10μm (B-C’, D-E’, M-M’’, N-N’’). The white dotted line indicates the border of the niche (B-C’, M-N’’), or the border of the LG (K-K’’, P-P’’). Statistical analysis was performed using Student’s t-test with Welch’s correction. Values are displayed as Mean ± SD. p-values indicating statistical significance have been mentioned in the respective graphs.

Next, we looked at Insulin signaling status in the LG in an HSD-induced insulin resistance model using the tGPH reporter. We observed that both in the Antp-positive niche and the progenitors flanking the dorsal vessel, the tGPH is localized more in the cytoplasm than the membrane, indicating low insulin sensitivity in the HSD-fed larval LG as compared to the CD-fed control (Fig. 7B-F). This indicates that HSD indeed affects the insulin signaling in the LG niche and MZ progenitors as well. Then, we checked blood cell homeostasis and studied LGs from larvae where we depleted ImpL2 in the hemocytes (*hml>UAS-ImpL2RNAi*) and fed them with HSD. We observed that in the HSD-fed insulin resistance scenario, when we stain the LGs with Antp, we indeed observe that the Antp-positive niche cell numbers decrease as compared to CD-fed larvae very similar to the LGs from tumor-bearing larvae, however, we observe that the niche numbers are rescued in the HSD-fed larvae where ImpL2 is depleted from hemocytes which provides an important regulatory role for hemocyte-derived ImpL2 on LG blood cell homeostasis (Fig. 7G-H). In corroboration with earlier results, when stained for plasmatocytes, we observe that there is an aberrant plasmatocyte differentiation in HSD-fed larvae as compared to CD-fed larvae; however, we observe that the increase in plasmatocyte numbers is rescued in the HSD-fed larvae, where ImpL2 is depleted from hemocytes (Fig. 7I-J). Additionally, when visualized with progenitor marker DE-cadherin, the progenitor pool that goes down in the HSD-fed larvae is rescued in HSD-fed larvae where ImpL2 is depleted in the Hml-positive hemocytes (Fig.7K-L). To find out whether this rescue of the niche phenotype is insulin signaling dependent or not, we stained the LG niche with pAkt and co-stained with Antp and found out that pAkt levels, a readout of insulin signaling are downregulated in the niche in the HSD-fed larvae as compared to CD-fed larvae and the pAkt levels are rescued in HSD-fed larvae where ImpL2 is depleted in the hemocytes (Fig.7M-O). Altogether, these results indicate that the hemocyte-derived ImpL2 regulates LG homeostasis in an insulin resistance scenario. Lastly, we observed that Wg expression goes down in the progenitors in the HSD-fed conditions as compared to the CD control, and this is rescued when ImpL2 is depleted in the hemocytes in HSD conditions (Fig. 7P-Q). Altogether, these results imply that a systemic metabolic stress like insulin resistance is indeed sensed by the LG for the maintenance of a fine balance between progenitor maintenance and differentiation. We also identify that circulating hemocyte-derived ImpL2 plays a regulatory role in controlling blood cell homeostasis both in cancer-induced cachexia and high sugar diet-induced insulin resistance scenario.

To determine the clinical relevance of our study, we examined IGFBP7, (the mammalian homologue of ImpL2) expression pattern in patient datasets. Transcriptomics analysis of TCGA data showed that *IGFBP7* mRNA levels are significantly upregulated in colon adenocarcinoma (n= 286) compared to normal colon tissue (n = 41) (Fig. S8A), which was corroborated by CPTAC proteomic data analysis (Fig. S8B). Critically, patients with higher IGFBP7 expression exhibited poorer overall survival (Fig. S8C). Furthermore, cell-type-specific analysis revealed elevated IGFBP7 expression in tumor-associated monocytes/macrophages compared to normal tissue macrophages (Fig. S8D), implicating IGFBP7’s role in immune cell dysregulation during colon cancer progression. A mechanistic correlation of IGFBP7 and colorectal cancer-induced cachexia needs further exploration.

Overall, our study establishes a model where we show that *yki^3SA^* larval gut tumors cause cachexia with characteristic hallmarks. We show that a known cachectic ligand, ImpL2, is not just upregulated in the tumor but also in the circulating hemocytes. Our study unravels a critical role of ImpL2 in regulating LG blood cell homeostasis in a tumor-induced insulin resistance condition. Elevated ImpL2 levels cause an abrogation of insulin signaling in the niche-progenitor micro-environment, impacting progenitor maintenance, pushing them towards differentiation (Fig. 8).

**Fig 8:**
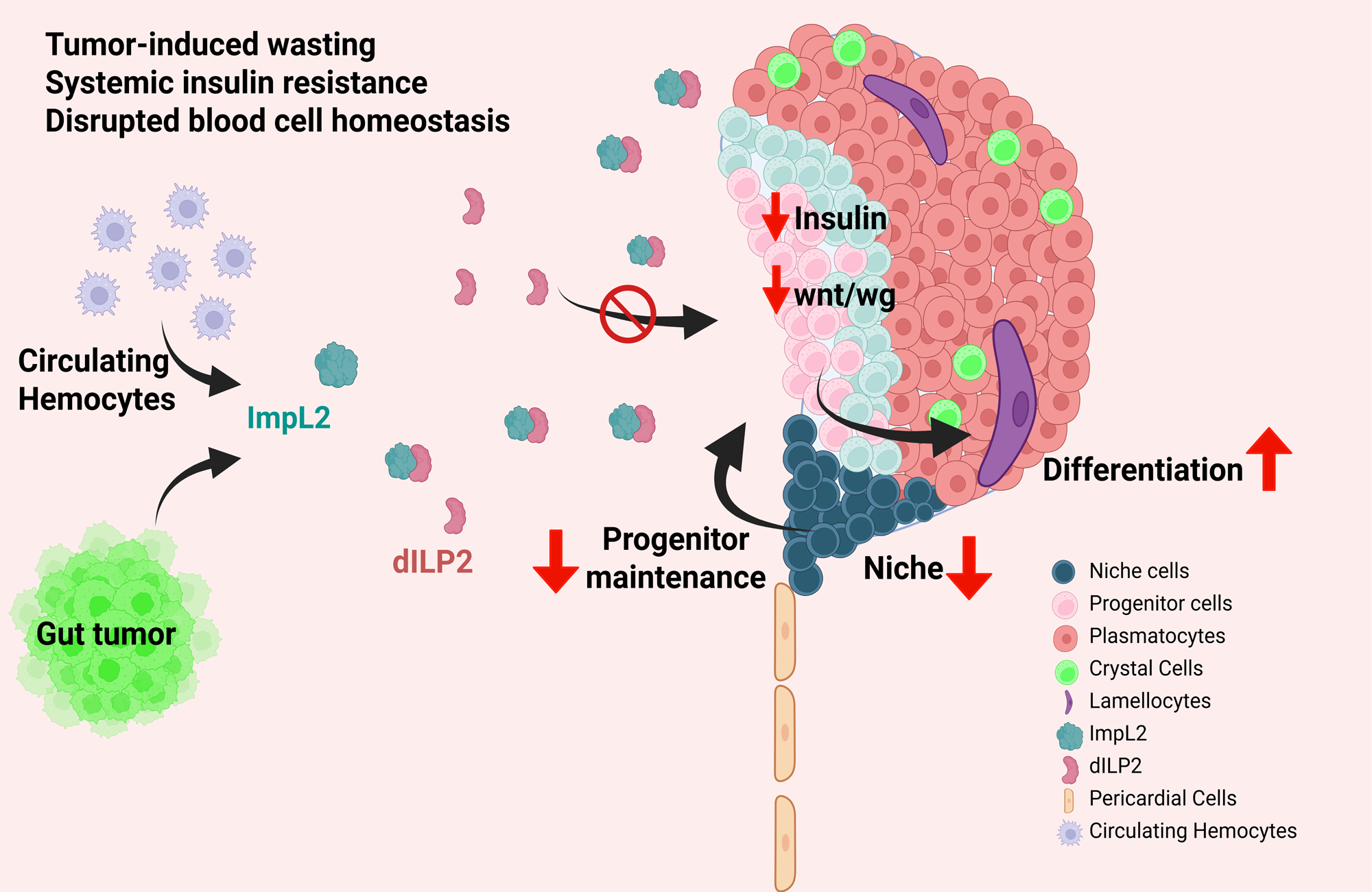
Graphical abstract depicting tumor-induced systemic insulin resistance perturbs blood cell homeostasis in the *Drosophila* model of cancer cachexia. A mechanistic model illustrating how a tumor induces systemic insulin resistance and disrupts peripheral tissue homeostasis. The tumor triggers a systemic increase in the insulin antagonist ImpL2, which binds to and inhibits the function of the insulin-like peptide, Dilp2. This results in profound consequences for the hematopoietic system, disrupting lymph gland homeostasis and leading to a smaller niche and aberrant differentiation of progenitors into mature blood cells. The model also highlights a crucial role for hemocyte-derived ImpL2 in exacerbating this systemic disruption of tissue homeostasis.

## Discussion

Cancer cachexia has long been recognized as a consequence of tumor-host interactions and systemic inflammation. However, the mechanisms linking tumors to hematopoietic dysfunction, immune dysregulation, and subsequent tissue wasting remain largely unknown^28,86^. Mammalian studies have shown that indeed epithelial tumors disrupt hematopoiesis^27,28,87^, and there is emerging evidence suggesting that altered blood cell homeostasis can be an active driver of tumor-associated systemic inflammation^29^. In cancer patients, bone marrow exhibits increased myelopoiesis with preferential differentiation towards inflammatory cell lineages, a phenomenon termed “myeloid skewing” that correlates with cachexia severity and poor prognosis^29,88^. Tumor-derived factors and systemic inflammation drive alterations in hematopoiesis, leading to emergency myelopoiesis, skewed differentiation programs, and an immunosuppressive environment, which results in tumor progression^29,89,90^. We demonstrate that in the *Drosophila* larval cachexia model, the lymph gland (LG), analogous to mammalian bone marrow, has a reduced niche size and aberrant blood cell differentiation. Our study represents an important first step in characterizing how blood cell homeostasis is affected in cancer cachexia. One of the main questions that emerges from our study is whether these LG-derived differentiated hemocytes act as passive bystanders merely reflecting systemic disease, or do they actively contribute to cachexia progression through inflammatory signaling or by rewiring the overall metabolic state, thereby impacting multiple organs.

Larval hemolymph consists of circulating hemocytes that arise from the first wave of hematopoiesis that occurs in the embryo^35,37,91^. Recent single-cell sequencing studies have shown that the three different blood cell types, i.e., plasmatocytes, crystal cells, and lamellocytes, are very heterogeneous and belong to more than one subtype^39,50^.The role of hemocytes has been promiscuous when it comes to epithelial tumors, some showing that circulating hemocytes are recruited to the tumors and result in tumor-growth retarding effects, and on the contrary, others suggesting upregulation of factors that promote tumor growth^55–59,92^. Apart from the role of circulating hemocytes in regulating tumor growth, hemocyte function in conferring systemic inflammation, particularly in cancer cachexia holds immense significance. However, apart from a few studies suggesting that immune-and coagulation-related factors are heightened in cancer cachexia^93,94^, the gene signature of circulating hemocytes in cachexia is largely unknown. We have performed bulk transcriptomics analyses of circulating hemocytes in cachectic larvae, which suggests upregulation of multiple known cachectic ligands, like ImpL2, Pvfs, and inflammatory ligands like Spaetzle, etc. It was earlier shown in adult cachexia models that apart from the tumor, distant organs like muscle or fat body also have higher ImpL2 levels^31,32,61,62,69,78^; however, we report for the first time that circulating hemocytes have elevated ImpL2 levels even in the circulating hemocytes in a larval cachexia model. Our study highlights the role of ImpL2 in the dysregulation of LG blood cell homeostasis in cachexia. Our observations indicate that there is systemic insulin resistance and hyperglycemia in the larval cachexia model, as shown earlier in various cachexia models^31,61,62^. However, it needs to be noted here that ImpL2 might not be the sole contributor to systemic insulin resistance, and there could be other signaling components that cause a resultant insulin resistance condition. Our findings suggest that *yki^3SA^*-induced larval gut tumor results in systemic insulin resistance typical of cancer patients^15,16,70,95^. LG responds to systemic metabolic stress like starvation and infection^42,96^; however, how the LG senses cues from a distant epithelial tumor is completely unexplored. This study looks into a fairly under-characterized aspect of deciphering whether the blood cell homeostasis is affected during cancer cachexia. LG progenitors integrate systemic metabolic cues to guide lineage commitment decisions^38^, and examining the hematopoietic system offers unique insights as blood cells might serve as both targets and perpetrators of metabolic dysfunction. ImpL2 derived from the tumor has already been shown to aggravate and result in peripheral tissue wasting and systemic insulin resistance^31,32,62^. Here, we show that elevated ImpL2 caused by the tumor, circulating hemocytes, results in disruption of hematopoiesis.

Insulin signaling plays a critical role in developmental hematopoiesis in niche and progenitor maintenance in the LG^42,43,77^. However, in a condition like tumor-induced systemic insulin resistance, where there is a low insulin sensitivity in most peripheral organs, it is completely unclear as to how the LG responds. In this study, we investigated the status of Insulin signaling in the LG niche-progenitor micro-environment, and we find that Insulin signaling is inactive both in the niche and the progenitors in the cachexia background. These observations imply that systemic insulin resistance abrogates the ability of LG progenitors to sense systemic signals that activate insulin signaling in the MZ for progenitor maintenance. It would be worth exploring if the Insulin activation status is similar in the circulating hemocytes and whether this has any bearing on their ability to be either pro- or anti-tumorigenic^97^, as has been shown for mammalian macrophages, where PI3K-Akt signaling dictates whether the macrophages would be M1 or M2^98–100^. Here, we show that elevated ImpL2 levels in a tumor-induced cachexia background or during HSD conditions affect LG homeostasis by impacting Insulin signaling. Now, an earlier study states that Insulin signaling in the MZ acts upstream of Wg signaling in controlling progenitor fate^42^. However, there are multiple signaling pathways, like JAK-STAT, Dpp, Hedgehog, and Notch, that play a critical role in progenitor maintenance^36,101–106^.

We find that Wg signaling is perturbed in the LGs from cachectic larvae; however, it needs to be studied if Insulin signaling directly impacts Wg signaling functioning upstream in the cachexia scenario, or if this is a secondary effect where multiple progenitor maintenance pathways are impacted. Under HSD conditions, which we used as a mimic of the insulin resistance scenario caused by the tumor, we find that ImpL2 is upregulated even in the circulating hemocytes, causing a similar effect on LG hematopoiesis as in the case of cachexia, and the LG phenotypes can be rescued by depleting ImpL2 from hemocytes in HSD conditions. Interestingly, our data also shows that hemocyte-specific overexpression of ImpL2 alone is capable of mirroring the insulin resistance phenotype of down-regulation of pAkt levels in the fat bodies of larvae bearing hemocyte-specific ImpL2 overexpression. Chronic inflammation is central to the pathophysiology of Type 2 Diabetes Mellitus (T2DM) or Insulin Resistance (IR)^95,107^. IR profoundly impacts hematopoiesis through metabolic, inflammatory, and cellular mechanisms. IR-associated chronic low-grade inflammation promotes myeloid-biased differentiation^72,108,109^. Elevated IL-1β, TNF-α, and IL-6 in IR states drive emergency myelopoiesis^109^. However, the mechanistic underpinnings of tumor-induced insulin resistance, especially in conditions like cancer cachexia, have not yet been explored. In this context, the role of ImpL2 homolog IGFBP7 is underexplored. TCGA datasets of colorectal adenocarcinoma patients suggest that higher *IGFBP7* expression is associated with a poorer survival rate. Now, it should be noted here that the occurrence of cancer cachexia in colorectal cancer patients is high^110–112^. However, the specific role of IGFBP7 in cancer cachexia or hematopoietic-immune remodeling remains to be explored.

Our study shows that the abrogation of Insulin signaling leads to niche size reduction and aberrant prohemocyte differentiation into the mature blood cell types. A longitudinal developmental study would be needed to assess if these cells that go out in the adult circulation are functionally more active or not, as compared to their non-tumor background counterparts. An important aspect would be to investigate whether these circulating hemocytes foster tumor growth by supplying nutrients ferried in from wasted tissues, or are supporting the rejuvenation and repair of wasted tissues. Our study sheds light on a previously under characterized inter-organ crosstalk between the tumor and the hematopoietic system. This crosstalk would be important to understand the basis of systemic chronic inflammation and tumor-host interactions that could be mediated via blood cells. Our study opens up future avenues of investigation in understanding how the hematopoietic system transitions from being a target of tumor-induced stress to becoming a contributor to metabolic dysfunction.

## Materials and methods

All the materials and methods have been described in the Supplementary Information

## Acknowledgements

We would like to thank the Digital Imaging Facility (DIF) and the Common Instrumentation Facilities (CIF) at ACTREC for all the support. We thank the Bloomington *Drosophila* Stock Center, Developmental Studies Hybridoma Bank, and the fly community for fly stocks and antibodies. We would like to particularly thank Thomas Dolezal, Hugo Stocker, Lolitika Mandal, Gaurav Das, and Tina Mukherjee for various fly lines and reagents. We would like to thank David Bilder, Tor Erik Rusten and Thomas Dolezal for insightful suggestions. We are thankful to the Stem Cell and Tissue Homeostasis lab for useful input and discussions.

## Author contributions

Conceptualization: R.J.K.; Data curation: R.J.K., U.C.; Formal analysis: U.C., G.P., P.K., P.B. and T.T.; Funding acquisition: R.J.K.; Investigation: U.C., G.P., P.K., P.B., T.T.; Methodology: U.C., G.P., P.K., P.B., T.T.; Project administration: R.J.K.; Resources: R.J.K.; Supervision: R.J.K.; Validation: U.C.; Visualization: R.J.K.; Writing – original draft: U.C., R.J.K.; Writing – review & editing: U.C., R.J.K.

## Data availability

All relevant data can be found within the article and its supplementary information

