## Supplementary Information for "*Drosophila* midgut tumor-induced insulin resistance systemically remodels lymph gland hematopoiesis during cancer cachexia" (from another version of this preprint: version 1, 2025-11-26)

**Materials and Methods:**

**Fly husbandry and strains:**

All fly stocks and genetic crosses were maintained on standard cornmeal medium at 25℃, 12:12 h light/dark cycles unless otherwise indicated. For High Sugar Diet (HSD) experiments, cultures were maintained on standard cornmeal medium containing sucrose, 0.15M for Control Diet (CD) and 1.0M for HSD, respectively, as the only purified sugar source[1]. For all *Gal4*-based analysis, respective *Gal4* lines were crossed to a wild-type genetic background and were used as controls. Canton-S was used as the wild type. For analysis of mutant alleles, the wild type was used as a control. Fly stocks used were as follows: *esg-Gal4, UAS-GFP/CyO*, *UAS-yki$^{3SA}$* (RRID: BDSC_28817), *UAS-ImpL2-RA/TM3Sb, UAS-s.ImpL2*, *UAS-ImpL2RNAi* (RRID: VDRC_KK106543) (gift from Dr. Hugo Stocker, ETH Zurich), *hml-Gal4GFP, fb-Gal4*, *domeMESO-Gal4GFP* (gift from Dr. Tina Mukherjee, inStem, Bangalore), *tepIVGal4- GFP, antp-Gal4.UAS-mCD8-RFP/Tb* (gift from Dr. Lolitika Mandal, IISER Mohali), *Df(3L)Ilp2-3,Ilp5$^{3}$* (RRID: BDSC_30888), *Df(3L)Ilp2-3* (RRID: BDSC_30889), *UAS-ImpL2RNAi* (RRID:BDSC_64936), *UAS-dilp2RNAi*, *dilp2-Gal4/CyOGFP* (gift from Dr. Gaurav Das, NCCS, Pune),*tGPH* (RRID: BDSC_8164)

**Antibodies:**

Antibodies used were mouse anti-Cora (1:25, C566.9, DSHB; Ab_1161642), mouse anti-Cora (1:25, C615.16, DSHB; Ab_1161644), mouse anti-P1 (1:100, kind gift from Dr. Istvan Ando), mouse anti-Hindsight (1:25, 1G9 – DSHB; RRID: Ab 528278), mouse anti-Antp (1:25, 8C11-DSHB; RRID: Ab_528083), rat anti-DECAD (1:25, DSHB DCAD2; RRID: Ab_528120) mouse anti-Myospheroid (1:25, CF.6G11 - DSHB; RRID: Ab_528310), mouse anti-Wg (1:25, 4D4-s-DSHB; RRID: AB_528512), rabbit anti-pAkt (1:100, 4060-S Cell Signaling Technologies, RRID: AB_2315049), rat anti-dilp2 (1: 250, gift from Dr. Pierre Leopold), mouse anti-phospho-Y489-β-catenin (pY489-β-cat) (1:100, DSHB; Cat# PY489-B-catenin; RRID: AB_10144551), for immunofluorescence based experiments. Normal Goat Serum (HIMEDIA, RM10701) was used as the blocking agent. Alexa-Fluor 568 and 488 conjugated secondary antibodies – Goat-raised anti-mouse 568 (1:400, Invitrogen, RRID: Ab_144696), and Goat-raised anti-mouse 488 (1:400, Invitrogen, RRID: AB_2534069), Goat-raised anti-rabbit 488 (1:400, Invitrogen, RRID: AB_10563566), Goat-raised anti-rat 568 (1:400, Invitrogen, RRID: AB_10563566), were used for immunofluorescence-based experiments.

**Dissections and Immunofluorescence:**

Larval guts were dissected in 1x Phosphate Buffer Saline (PBS), fixed in a 1:1 ratio of n-heptane and 4% paraformaldehyde (PFA), and incubated for 30 minutes at 29℃. After removing PFA, samples were washed with methanol for 10 seconds, followed by three PBS washes. Samples were then incubated in 1x PBS containing 0.1% Triton-X-100 (PBST) for 10 minutes. Blocking was performed using 20% normal goat serum for 20 minutes, followed by overnight incubation with respective primary antibodies at 4℃. This was followed by 1x PBS washes, blocking, and incubation with appropriate Alexa-Fluor conjugated secondary antibody for two hours at room temperature, then three washes with 1x PBS for 5 minutes each. The guts were then mounted in Vectashield Antifade mounting medium containing DAPI (Vector Laboratories, RRID: AB_2336790).

Larval fat bodies, lymph glands (LGs), and brains (to stain the Insulin Producing Cells or IPCs) were dissected in phosphate buffer saline (PBS), fixed in 4% paraformaldehyde for 20 minutes, followed by three washes with 1x PBS containing 0.3% Triton-X (PBST) for 5 minutes each. The samples were then blocked in 20% normal goat serum (HIMEDIA, RM10701) for 20 minutes at room temperature, followed by overnight primary antibody incubation at 4°C. This was followed by 0.3% PBST washes, blocking, and incubation with the appropriate Alexa-Fluor conjugated secondary antibody for 2 hours at room temperature, then three 0.3% PBST washes for 5 minutes each. The fat bodies, LGs, and brains were then mounted in Vectashield Antifade mounting medium containing DAPI and imaged.

**Nile Red staining and lipid droplet (LD) area quantification:**

Fat bodies from third instar larvae were dissected in PBS and immediately fixed in 4% paraformaldehyde for 45 minutes. They were washed thrice in 1x PBS and incubated for 40 minutes in diluted Nile Red Solution (1:200 from 200µg/ml stock in DMSO, Sigma N3013) at room temperature. Samples were washed again in PBS and mounted in Vectashield Antifade mounting medium containing DAPI and imaged the same day.

Lipid droplet measurements were acquired in a 40x oil objective at 1x zoom, and lipid droplet area measurements were performed as described in the study[2] . 4 ROIs per fat body and fat bodies from ten larvae per genotype were taken for analysis.

**2-NBDG uptake assay and quantification:**

2-NBDG uptake was performed in the fat body as described in the study[3]. Fat bodies were incubated in 2-NBDG (Invitrogen, N13195) diluted 1:100 in 1x PBS for 15 minutes in the dark, rinsed twice in 1X PBS, and fixed in 4% paraformaldehyde for 30 minutes at room temperature. Tissues were rinsed again, mounted in Vectashield Antifade mounting medium containing DAPI, and imaged the same day.

For measuring fluorescence intensity of 2-NBDG in the fat body samples, images were acquired using the same settings for both control and tumor genotypes, and the mean fluorescence intensity was recorded from the maximum intensity Z-projected images.

**Immunostaining of muscle fillets and analysis:**

Heat fixation of larvae was performed as described in the study[4]. Briefly, larvae were collected, rinsed in cold 1X PBS, and transferred to a cell strainer (Corning CLS431750). The larvae were dipped into 65-70℃ water for 2-4 seconds and allowed to cool in ice-cold 1X PBS. Larvae were then dissected as described in the study[2]. Muscle fillets were then incubated in Phalloidin Atto 488 (1:200, Sigma 49409) for 30 minutes at room temperature, washed in 1X PBS, and mounted in Vectashield Antifade mounting medium containing DAPI and imaged.

For muscle thinning measurements, images of each muscle fillet were acquired using a 40x oil objective at 2x zoom. The myofiber width and the inter-myofiber distance were measured using the Line Tool in ImageJ, and the length was measured and recorded. Three ROIs per muscle fillet with at least five cuticular muscle fillets per genotype were taken for analysis.

**qPCR and primers used:**

For each biological replicate, RNA was extracted from five whole larvae. For gut samples, a total of seven larval guts were dissected in cold 1X PBS. For hemolymph samples, 80-100 larvae were bled in 1X PBS to extract RNA. Samples were then lysed in 500µL of Trizol Reagent (Ambion–Life Technologies, 11596018). Total RNA was extracted using the phenol-chloroform method, and RNA yield was quantified using a Nanodrop spectrophotometer. 1µg of RNA was reverse transcribed using the RevertAid First Strand cDNA Synthesis Kit (Thermo Fisher Scientific, K1622). Quantitation of the transcripts was performed using SYBR green chemistry in the QuantStudio 5 RT PCR system (Thermo Fisher Scientific). Data were analyzed using the ΔΔCt method, and relative mRNA expression was normalized to rp49. Fold change was calculated relative to the wild-type control. The experiment was performed in three biological replicates. Primer sequences used in qPCR experiments are mentioned below.

**Primers written in 5' to 3' direction**

Imp-L2 FP AAGAGCCGTGGACCTGGTA

Imp-L2 RP TTGGTGAACTTGAGCCAGTCG

Dilp2 FP ATGAGCAAGCCTTTGTCCTTC

Dilp2 RP ACCTCGTTGAGCTTTTCACTG

rp49 FP AGCATACAGGCCCAAGATCG

rp49 RP TGTTGTCGATACCCTTGGGC

**Circulating trehalose measurement:**

For circulating trehalose measurements, 30 third instar larvae were bled per replicate, and 1.5 µL hemolymph was used for each condition. Hemolymph samples were diluted in 99 µL of trehalose buffer TB (5mM Tris pH 6.6, 137mM NaCl, and 2.7mM KCl) and heated for 5 minutes at 70℃. Circulating trehalose estimation was done as described in the study [5]. Each sample was divided into two 50µL aliquots. To one aliquot, 50µL of TB was added; to the other 50µL of trehalose buffer containing Trehalase enzyme TS (1mL TB + 3µL Porcine trehalase, Sigma, T8778-1UN) was added. Trehalose and glucose standards (1, 0.16, 0.08, 0.04, 0.02, and 0.01 mg/ml, Trehalose from Sigma 90208) were prepared similarly. Samples and standards were incubated at 37℃ overnight. The next day, 30µL of each one was transferred to individual wells of a 96-well plate, and 100µL of GO reagent (Sigma, GAGO-20) was added. The plate was sealed and incubated at 37℃ for 45-60 minutes. To stop the reaction, 100µL of sulfuric acid was added. Absorbance at 540 nm was detected using a TECAN microplate reader. Concentrations were calculated based on the equation given by a linear regression and the sample dilution. The experiment was performed in three biological replicates. Statistical significance was determined by an unpaired Student's t-test with Welch's correction.

**Triglyceride measurement:**

For triglyceride quantification, five whole larvae were taken in 1 mL of 5 % NP-40, heated to 95°C for 5 minutes, cooled, and centrifuged at 14,000xg for 2 minutes. Supernatants were diluted 1:10 in Assay Buffer. Triglyceride quantification was performed using the Triglyceride Quantification Kit (MAK266, Sigma-Aldrich) according to the manufacturer's protocol. Briefly, a 50μL sample or standard was added to a 96-well plate, followed by 2 μL of lipase, and incubated for 20 minutes at room temperature. Next, 50μL of reaction mixture (46 μL buffer + 2μL probe + 2 μL enzyme mix) was added and incubated for 60 minutes at room temperature, protected from light. Absorbance was read at 570 nm. The experiment was performed in three biological replicates. Statistical significance was determined by an unpaired Student's t-test with Welch's correction.

**Image Acquisition and Analysis:**

All images were acquired in Zeiss LSM 780, Nikon AX, or Zeiss LSM 980 microscopes and quantified using ImageJ or Imaris 10.2.0 Image Analysis Software (RRID: SCR_007370).

Whole-mount LG samples were imaged with a 20x objective. Quantitation of the area of the primary LG lobe was done using the 'Freehand selection tool' to mark the area of the primary lobe, and measurements were recorded.

For pAkt intensity measurements, images were acquired using the same settings for both control and tumor genotypes, and then the Mean Fluorescence Intensity was recorded from the middle stacks (middle optical sections) of the images of the fat bodies. At least ten larvae per genotype were taken.

For Coracle quantification, guts were acquired using a 63x oil objective at 1x zoom at the same settings for both tumor and control groups. Coracle fluorescence intensity at cell-cell junctions was quantified by measuring mean intensity values using the Line Tool in ImageJ (Five junctions per gut, two ROIs per midgut). At least ten guts per genotype were taken for analysis.

For lymph gland (LG) analysis, the Plasmatocyte Differentiation Index was estimated by measuring the percentile of P1-positive area divided by the total area of the primary lobe. The freehand selection tool was used for measuring the area of the plasmatocytes. For the quantitation of Antp, Hnt, and pY489-β-cat, the positive signals for respective markers were manually counted using the Multi-Point Tool. For quantitation of Mys, the LGs were categorized as positive or negative based on the presence or absence of lamellocytes, and the percentage of lamellocyte-positive LGs was estimated. The LG quantitation was done for individual primary LG lobes.

For tGPH measurements in LG, both ROIs in the niche (co-stained with Antp) and the progenitors (middle optical sections merged) were set and then tGPH was quantified using the line tool in ImageJ; a line was drawn around the membrane for each cell to measure the membranous intensity and a rectangular box was drawn in the cytoplasmic region to measure the cytoplasmic intensity. Both the values were taken to calculate the fold change of the membranous/cytoplasmic intensity. Two to three cells per niche or MZ of LG were taken for analysis.

For pAkt in the niche and progenitors, similar ROIs were taken and estimated by calculating mean fluorescence intensity using the ImageJ software. For Wg and DE-Cadherin in the progenitors, middle stacks (middle optical sections merged) were taken and estimated by calculating mean fluorescence intensity using the ImageJ software.

For Dilp2 quantification in the IPCs, ROIs were drawn using the Free-Hand Tool around the IPCs in the larval brain samples positive for Dilp2 signal, and the Mean Fluorescence Intensity of maximum intensity Z-projected images were recorded and analyzed. At least six larval brains per genotype were taken for analysis.

Images shown in the figures are either projections of sub-slices or maximum intensity Z-projections of all optical sections. Images were processed using ImageJ. The raw images were converted to TIFF format (in RGB mode) into single or merged channels as per requirement using ImageJ. TIFF files were then opened using Adobe Illustrator (Creative Cloud version) and were assembled as per the requirements of the figure panel using a black background on an Illustrator canvas (RGB mode). Final assembled figure panels were saved in TIFF format using LZW compression with a resolution of 600 dpi or higher.

**Esg+ Cluster Volume Analysis:**

Three-dimensional volume analysis of Esg+ clusters was performed using the Surfaces function in Imaris 10.2.0 Image Analysis Software (RRID: SCR_007370). Surface rendering was generated using absolute intensity thresholding, which was maintained constant across all samples. A size filter of [>10 $\mu m^3$] was applied to exclude the background signal. Total cluster volumes per gut were quantified by summing individual clusters, especially the fused esg+ clusters in the tumor genotypes. Ten to fifteen guts per genotype were analyzed. Statistical analysis was performed using Student's t-test with Welch's correction in GraphPad Prism. Data are presented as mean ± SD.

**Transcriptomics analysis:**

**RNA Extraction and Quality Assessment**

Total RNA was extracted from hemolymph samples isolated from a hundred larvae. Hemolymph isolation was performed according to the protocol as described in the study [6]. Total RNA was isolated using the QIAgen RNeasy Mini Kit (Cat no 74104) following the manufacturer's instructions. RNA quality and integrity were assessed using an Agilent Bioanalyzer to ensure samples met the criteria for downstream RNA sequencing applications.

**Library Preparation**

RNA sequencing library preparation and sequencing were performed by MedGenome Labs Ltd. RNA sequencing libraries were prepared using a modified NEBNext Ultra II Directional RNA Library Prep Kit protocol. Poly-A-containing mRNA molecules were selectively purified using oligo-dT attached magnetic beads. Following purification, the mRNA was fragmented using divalent cations at elevated temperature. First-strand cDNA synthesis was performed using reverse transcriptase and random hexamers. The cDNA was then converted to double-stranded cDNA with uracil incorporated in place of thymine to preserve strand specificity. Strand directionality was maintained through USER enzyme-based digestion of the second strand, leaving a single functional strand that maps to the original transcribed DNA strand.

The USER-digested single-strand molecules were enriched and indexed through limited-cycle PCR amplification, followed by AMPure bead purification to generate the final cDNA libraries for sequencing. Prepared libraries were sequenced on an Illumina HiSeq X/NovaSeq platform to generate 2×150 bp paired-end reads per sample. Sequencing quality was maintained with up to 75% of sequenced bases achieving Q30 quality scores. Raw sequencing data were processed to generate FASTQ files and uploaded to an FTP server for downstream analysis.

**Data analysis:**

Raw sequencing data were subjected to quality assessment to identify and remove low-quality reads, bases, and adapter sequences. Adapter and contaminant sequences, including tRNA and rRNA, were removed by mapping the reads to the human genome and filtering structural RNA-derived reads. High-quality, cleaned reads were aligned to the *Drosophila melanogaster* reference genome (dm6) using HISAT2. Alignment statistics were generated to assess data retention at each stage of quality filtering, contamination removal, and mapping. Gene expression levels were quantified using FeatureCounts to obtain raw read counts, which were subsequently normalized with DESeq2. Differential gene expression analysis was performed using DESeq2, and differentially expressed genes with statistical significance were identified and reported. Adjusted p-value was taken to be lower than or equal to 0.05. A subsequent search of transcriptomic data for enhanced and silenced pathways and biological processes was done by using WEBGEStalt. PathON [7] was used to map target genes of 14 canonical signaling pathways and ligands, and plotted from the dataset into heatmaps.

**Statistical analysis:**

For immunofluorescence experiments of larval gut tumors and fat bodies, at least ten guts or fat bodies from ten larvae per genotype were analyzed. For hematopoietic parameters, at least ten LGs were analyzed per genotype (N=10), with each data point representing one primary LG lobe (n=20). For tGPH quantitation in the LG, each data point represents one cell, with two to three cells quantified per niche or MZ (n=30). For brain samples, at least six larval brains were used (N=6), with each data point representing the IPCs from one lobe of the brain (n=12). For muscle fillet, at least five cuticular muscle fillets were used (N=5), with each data point representing values from measurements of myofiber width and inter-myofiber distance from three ROIs per fillet (n=30).

Statistical analysis was performed using the GraphPad Prism Version 8.0.1 software (RRID: SCR_002798). For all experiments, each experimental sample was compared to its respective control. Mutant genotypes were compared to the wild-type controls, and the knockdown or overexpression genotypes were compared to their respective parental Gal4 controls that were crossed to wild-type. Statistical significance was determined by using a two-tailed unpaired Student's t-test with Welch's correction. p-values are indicated in the respective graphs. Data are represented as Mean ± SD. No statistical method was used to pre-determine the sample size, and the experiments were not randomized.

AI was used to proofread parts of the text and correction of grammar. The authors however reviewed and edited the content as needed.

**Supplementary Figure legends:**

**Fig S1: *yki$^{3SA}$*-induced larval gut tumor causes bloating and an increase in lipid droplet area**

Larval phenotype upon *yki$^{3SA}$* expression driven by *esg-Gal4, UAS-GFP*, as compared to the wild type control (A). Black arrows indicate bloating. (B) Heatmap representing lipid droplet (LD) area distribution quantified as surface area (µm²) in larvae bearing tumor (*yki$^{3SA}$*) compared to the wild type control. *esg-Gal4::UAS-GFP* crossed to *Canton-S* was used as the wild type. Scale Bar: 100µm (A).

**Fig S2: Larvae bearing *yki$^{3SA}$* gut tumors induce crystal cell and lamellocyte differentiation in the LG**

Whole-mount larval LG displaying primary, secondary, and tertiary lobes of larvae bearing tumor (*yki$^{3SA}$*) compared to the wild type control (A-A'). Graph showing the quantitation of the area of the primary LG lobe of larvae bearing tumor (*yki$^{3SA}$*) compared to the wild type control (B). LG stained with Hnt (Magenta; C-D) or Mys (Magenta) of larvae bearing tumor (*yki$^{3SA}$*) compared to the wild type control (F-G). Corresponding graph showing the quantitation of the number of Hnt-positive crystal cells (E) or the percentage of lamellocyte-positive LGs (H). *esg-Gal4::UAS-GFP* crossed to *Canton-S* was used as the wild type. Nuclei were stained with DAPI (Grayscale; Blue). Scale Bar: 200µm (A-A'), 50µm (C-D; F-G). Statistical analysis was performed using Student's t-test with Welch's correction. Values are displayed as Mean±SD. p-values indicating statistical significance have been mentioned in the respective graphs.

**Fig S3: Bulk mRNA sequencing of circulating hemocytes in the larval cachectic model suggests differential gene expression across various signaling pathways.**

Heatmap-based visualization of all differentially expressed genes in the bulk mRNA sequencing of circulating hemocytes of the larvae-bearing tumor (*yki$^{3SA}$*) compared to the wild type control, and corresponding Venn diagram showing the number of upregulated and downregulated genes in the bulk mRNA sequencing data (A-B). Heatmap colour intensity displays log2-fold change values relative to the mean gene expression level across both genotypes. Volcano Plot visualization of all differentially expressed genes (C) or differentially expressed ligands of canonical signaling pathways (D). Bar charts showing a list of Biological Processes, Cellular Components, and Molecular Functions enriched in the differentially expressed gene sets in the RNA-seq data. (E). Dot plot visualization of Gene Set Enrichment Analysis showing activated and suppressed pathways in the RNA-seq data (F).

**Fig S4: *dilp2* mutants exert perturbation in blood cell homeostasis.**

LG stained with Antp (Magenta) or P1 (Magenta) in *dilp-2,3,5* mutant larva (*Df(3L)Ilp2-3, Ilp5$^{3}$*) (B, F) and *dilp-2,3* mutant larva (*Df(dilp2-3)*) (C, G) compared to the wild type control (A, E).Corresponding graph showing the quantification of the number of Antp-positive niche cells (D) or Plasmatocyte differentiation index (H). LG stained with P1 (Magenta) upon *dilp2-Gal4* driven overexpression of ImpL2 (*dilp2>UAS-ImpL2*) (J) or knockdown of Dilp2 (*dilp2>UAS-dilp2$^{RNAi}$*) (K) compared to the wild type control (I). Corresponding graph showing the quantitation of the Plasmatocyte differentiation index (L). *Canton-S* was used as the wild type (A, E), *dilp2-Gal4>wild type* was used as the wild type (I). Nuclei were stained with DAPI (Blue). Scale Bar: 50µm (A-C, E-G, I-K). Statistical analysis was performed using Student's t-test with Welch's correction. Values are displayed as Mean±SD. p-values indicating statistical significance have been mentioned in the respective graphs.

**Fig S5: Localized perturbation of ImpL2 in the niche or progenitors alters LG blood cell homeostasis**

LG stained with P1 (Magenta) upon core-progenitor specific overexpression of ImpL2 (*tepIV>UAS-ImpL2*) (B) or pan-progenitor specific overexpression of ImpL2 (*domeMESO>UAS-ImpL2*) (E) compared to respective wild type controls; *tepIV>wt* (A) or *domeMESO>wt* (D). Corresponding graph showing the quantitation of the Plasmatocyte differentiation index of the respective groups (C, F). LG stained with Antp (Green) upon niche-specific knockdown of ImpL2 (*ImpL2$^{RNAi}$*) using *antpmRFP-Gal4* (Red) (H) compared to the wild type control (*antpmRFP>wt*) (G) and corresponding graph showing the quantification of the number of Antp-positive niche cells (I). Nuclei were stained with DAPI (Blue). Scale Bar: 50µm (A-H). Statistical analysis was performed using Student's t-test with Welch's correction. Values are displayed as Mean±SD. p-values indicating statistical significance have been mentioned in the respective graphs.

**Fig S6: ImpL2 depletion from the gut tumor does not rescue tumor growth but rescues cachexia phenotypes.**

Estimation of larval midgut-specific GFP-positive *esg* clusters (Green) upon depletion of ImpL2 in the genetic background of expression of *yki$^{3SA}$* using *esg-Gal4, UAS-GFP* (C), as compared to the *yki$^{3SA}$* induction using *esg-Gal4, UAS-GFP* tumor control (B) or the *UAS- Impl2$^{RNAi}$* driven by *esg-Gal4, UAS-GFP* alone control (A). Corresponding graph showing the quantitation of the total GFP positive *esg* cluster volume per gut (D). Larval fat bodies stained with Nile Red (Red) upon depletion of ImpL2 in the genetic background of expression of *yki$^{3SA}$* using *esg-Gal4, UAS-GFP* (G), as compared to the *yki$^{3SA}$* induction using *esg-Gal4, UAS-GFP* tumor control (F) or the *UAS-Impl2$^{RNAi}$* driven by *esg-Gal4, UAS-GFP* alone control (E) Corresponding graph showing the quantitation of the LD area (H). Muscle fillet stained with Phalloidin (Green) upon depletion of ImpL2 in the genetic background of expression of *yki$^{3SA}$* using *esg-Gal4, UAS-GFP* (K), as compared to the *yki$^{3SA}$* induction using *esg-Gal4, UAS-GFP* tumor control (J) or the *UAS-Impl2$^{RNAi}$* driven by *esg-Gal4, UAS-GFP* alone control (I). Corresponding graph showing quantitation of the myofiber width (L). Nuclei were stained with DAPI (Blue). Scale Bar: 50µm (A-C), 20µm (E-G, I-K). White dotted lines indicate the border of the midgut. Statistical analysis was performed using Student's t-test with Welch's correction. Values are displayed as Mean±SD. p-values indicating statistical significance have been mentioned in the respective graphs.

**Fig S7: ImpL2 is upregulated in High Sugar Diet (HSD)-induced insulin resistance conditions and also drives insulin resistance.**

Larval fat bodies stained with pAkt (Green) from HSD-fed larvae with hemocyte-driven (*hml-Gal4*, *UAS-GFP*; Green) knockdown of ImpL2 (*UAS-ImpL2$^{RNAi}$*) (C) and Control Diet (CD)- fed larvae with hemocyte-driven overexpression of ImpL2 (*UAS-ImpL2*) (D) compared to HSD-fed larvae (B) and CD-fed larvae (A). Corresponding graph showing the quantitation of the Mean Fluorescence Intensity of pAkt in the respective groups (E). IPCs of larval brains stained with Dilp2 (Red) from HSD-fed larvae with hemocyte-driven (*hml-Gal4*, *UAS-GFP*; Green) knockdown of ImpL2 (*UAS-ImpL2$^{RNAi}$*) (H) and CD-fed larvae with hemocyte-driven overexpression of ImpL2 (*UAS-ImpL2*) (I) compared to HSD-fed larvae (G) and CD-fed larval control (F). Corresponding graph showing the quantitation of the Mean Fluorescence Intensity of Dilp2 in the respective groups (J). *Canton-S* was used as the wild type. Scale Bar: 20μm (A-D, F-I). White dotted lines indicate the border of the IPCs in the larval brain. Statistical analysis was performed using Student's t-test with Welch's correction. Values are displayed as Mean±SD. p-values indicating statistical significance have been mentioned in the respective graphs.

**Fig S8. High IGFBP7 Expression in Both Tumor and Immune Cells Correlates with Adverse Survival in Colorectal Cancer**

Transcript per million (TPM) values for IGFBP7 in colon adenocarcinoma (COAD) tumor samples (n = 286) compared to normal colon tissue samples (n = 41) obtained from The Cancer Genome Atlas (TCGA) database. Statistical significance was assessed using the Mann-Whitney U test (A). IGFBP7 protein abundance in CPTAC colon cancer (n = 97) versus normal adjacent tissue (n = 100). Z-values represent standard deviations from the median across samples for the given cancer type. Log2 Spectral count ratio values from CPTAC were first normalized within each sample profile, then normalized across samples (B). Kaplan-Meier survival curves displaying the estimated survival probabilities of Colon Adenocarcinoma patients with low (Blue) and high (Red) expression of IGFBP7 based on data from the Pan Cancer Atlas of The Cancer Genome Atlas (TCGA) (C) Box plot showing IGFBP7 mRNA expression in Monocytes/Macrophages from colon adenocarcinoma (COAD) tumor samples (Blue) versus normal colon tissue (Orange). Data were analyzed using GEPIA2021 based on the TCGA and GTEx databases. The box represents the interquartile range (IQR), and the line inside the box indicates the median. (D). p-values have been mentioned in the respective graphs.

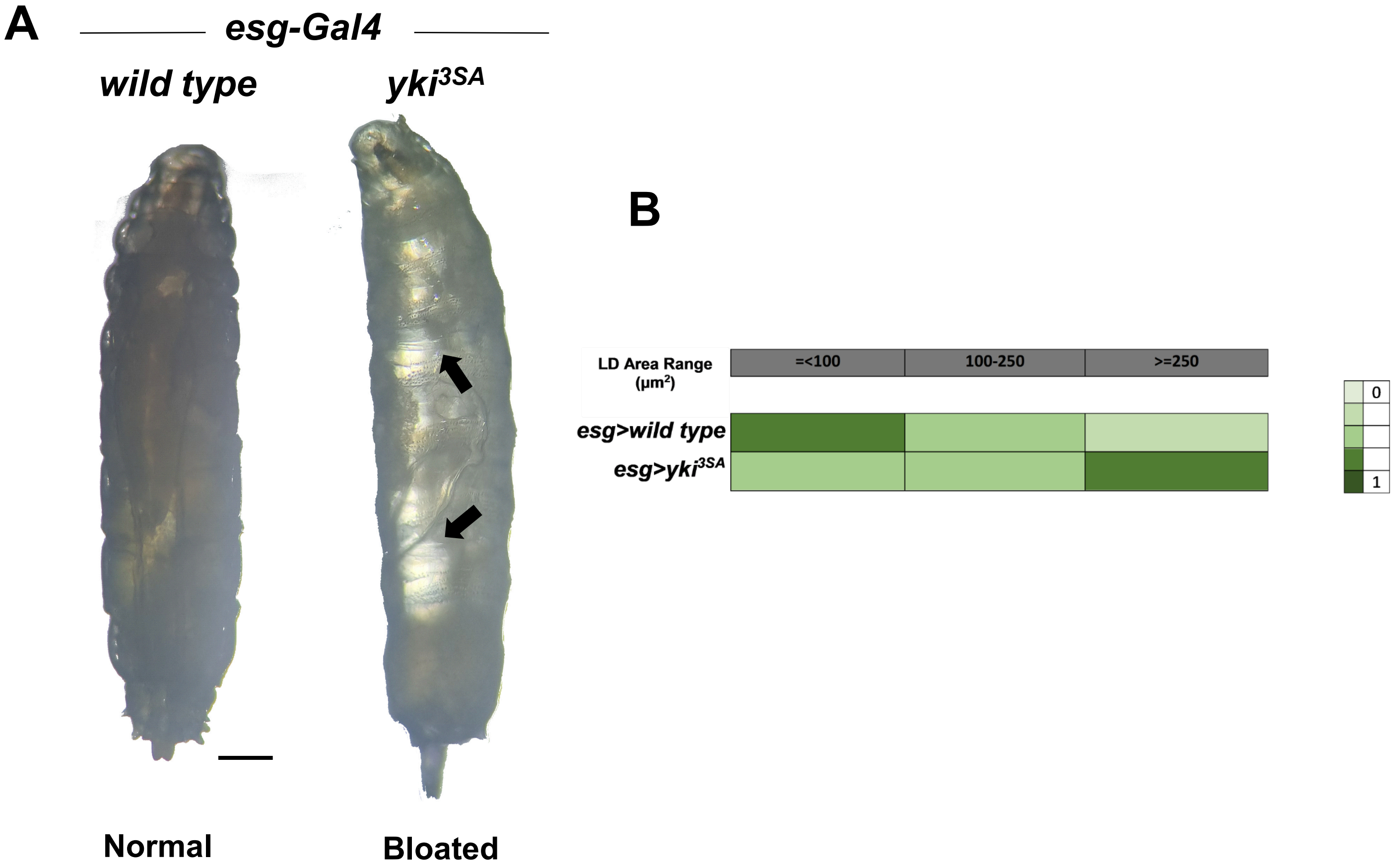

A
esg-Gal4
wild type
yki3SA
Normal
Bloated
B
LD Area Range (µm2)
=<100
100-250
>=250
esg>wild type
esg>yki3SA
0
1

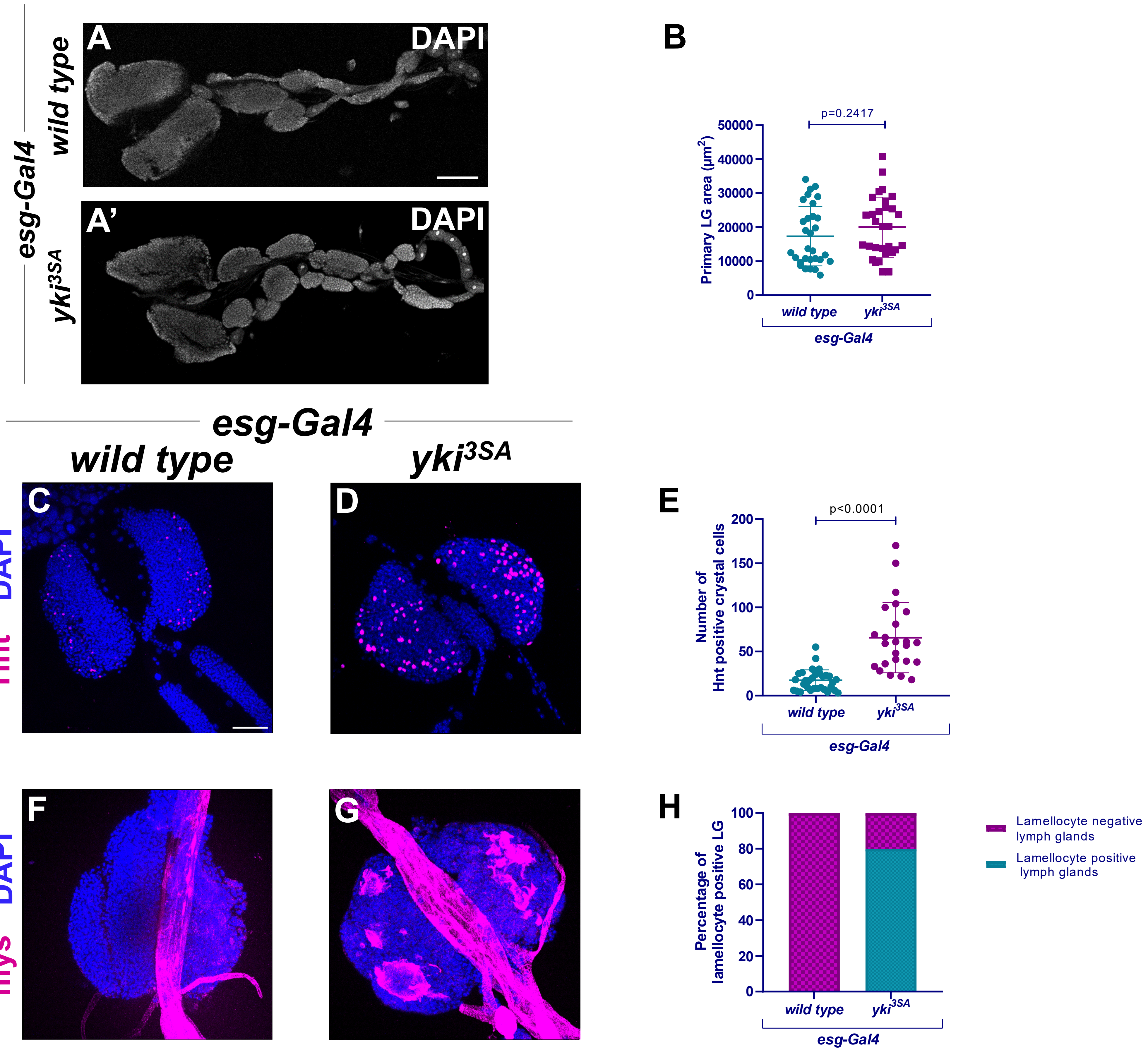
esg-Gal4
wild type
yki3SA
A
DAPI
A'
DAPI
B
p=0.2417
Primary LG area (μm2)
50000
40000
30000
20000
10000
0
wild type
yki3SA
esg-Gal4
esg-Gal4
wild type
yki3SA
C
D
Hnt DAPI
E
p<0.0001
Number of Hnt positive crystal cells
200
150
100
50
0
wild type
yki3SA
esg-Gal4
F
G
mys DAPI
H
Percentage of lamellocyte positive LG
100
80
60
40
20
0
wild type
yki3SA
esg-Gal4
Lamellocyte negative lymph glands
Lamellocyte positive lymph glands

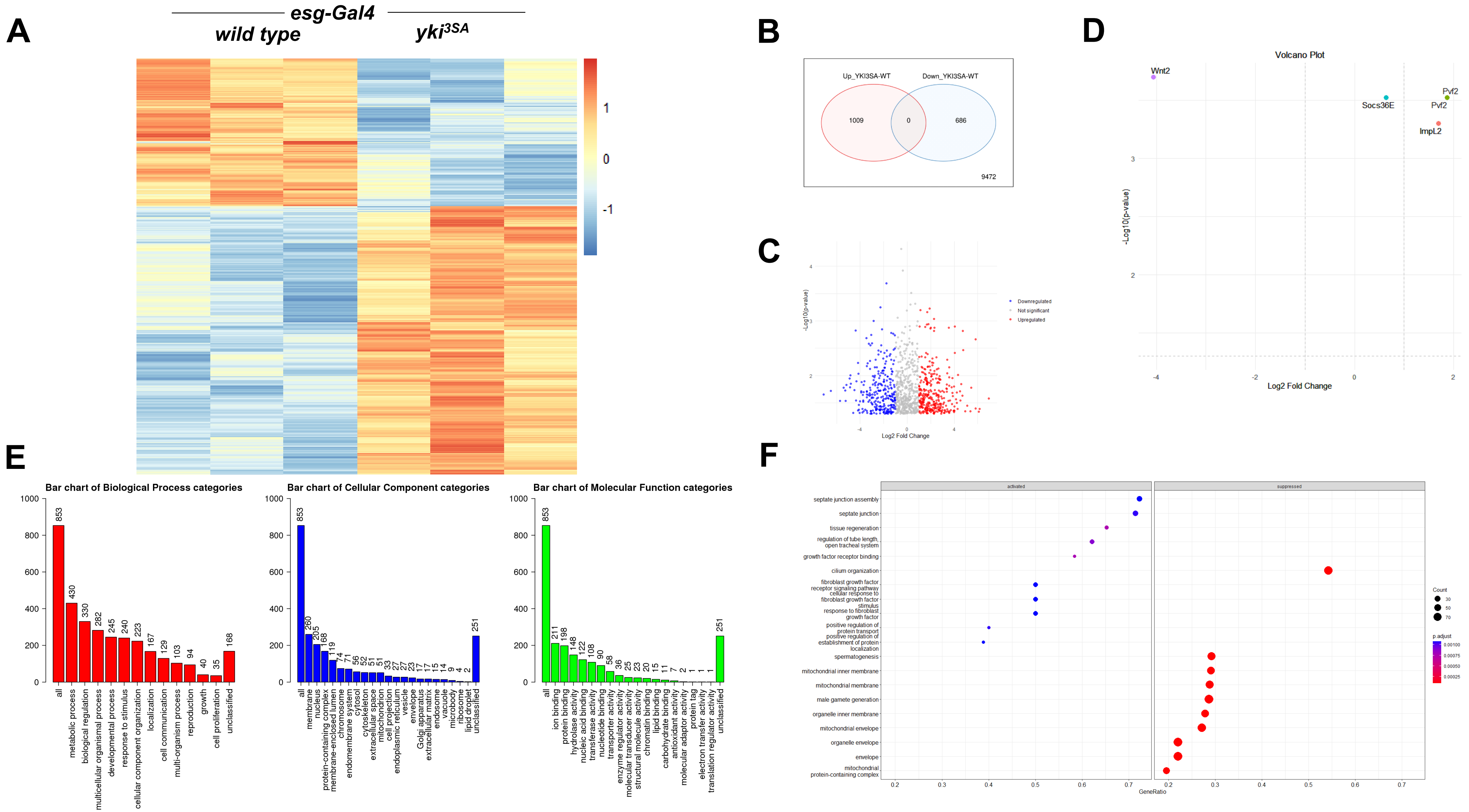

A
esg-Gal4
wild type
yki3SA
1
0
-1
B
Up_YKI3SA-WT
Down_YKI3SA-WT
1009
0
686
9472
C
Downregulated
Not significant
Upregulated
-Log10(p-value)
Log2 Fold Change
D
Volcano Plot
Wnt2
Socs36E
Pvf2
Pvf2
ImpL2
-Log10(p-value)
Log2 Fold Change
E
Bar chart of Biological Process categories
Bar chart of Cellular Component categories
Bar chart of Molecular Function categories
all
metabolic process
biological regulation
multicellular organismal process
developmental process
response to stimulus
cellular component organization
localization
cell communication
multi-organism process
reproduction
growth
cell proliferation
unclassified
membrane
nucleus
protein-containing complex
membrane-enclosed lumen
chromosome
endomembrane system
cytosol
cytoskeleton
extracellular space
mitochondrion
cell projection
endoplasmic reticulum
vesicle
envelope
Golgi apparatus
extracellular matrix
endosome
vacuole
microbody
ribosome
lipid droplet
ion binding
protein binding
hydrolase activity
nucleic acid binding
transferase activity
nucleotide binding
transporter activity
enzyme regulator activity
molecular transducer activity
structural molecule activity
chromatin binding
lipid binding
carbohydrate binding
antioxidant activity
molecular adaptor activity
protein tag
electron transfer activity
translation regulator activity
F
activated
suppressed
septate junction assembly
septate junction
tissue regeneration
regulation of tube length, open tracheal system
growth factor receptor binding
cilium organization
fibroblast growth factor receptor signaling pathway
cellular response to fibroblast growth factor stimulus
response to fibroblast growth factor
positive regulation of protein transport
positive regulation of establishment of protein localization
spermatogenesis
mitochondrial inner membrane
mitochondrial membrane
male gamete generation
organelle inner membrane
mitochondrial envelope
organelle envelope
envelope
mitochondrial protein-containing complex
GeneRatio
Count
p.adjust

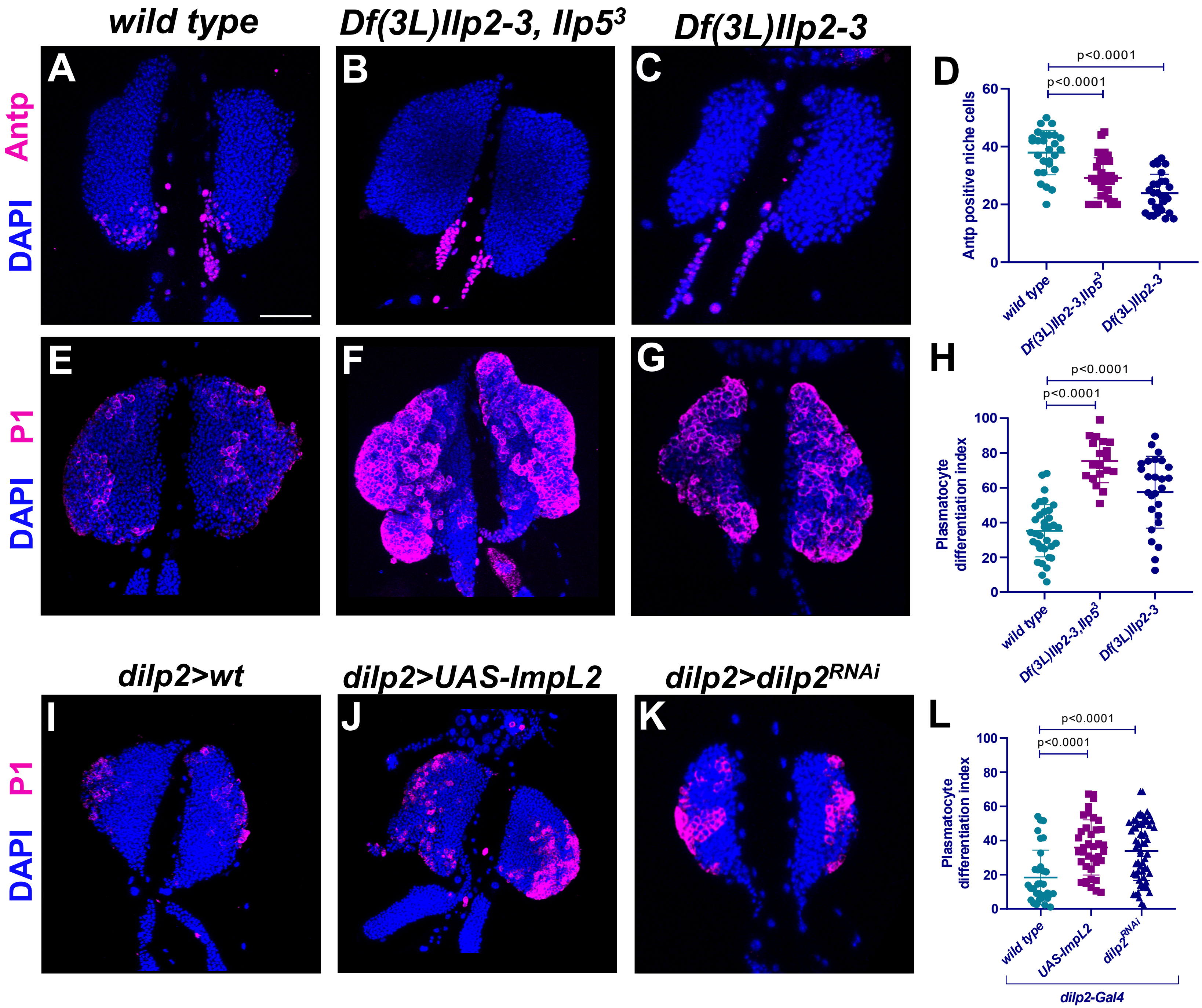

wild type
Df(3L)Ilp2-3, Ilp5³
Df(3L)Ilp2-3
A
B
C
D
Antp
DAPI
p<0.0001
p<0.0001
Antp positive niche cells
60
40
20
0
wild type
Df(3L)Ilp2-3,Ilp5³
Df(3L)Ilp2-3
E
F
G
H
P1
DAPI
p<0.0001
p<0.0001
Plasmatocyte differentiation index
100
80
60
40
20
0
wild type
Df(3L)Ilp2-3,Ilp5³
Df(3L)Ilp2-3
dilp2>wt
dilp2>UAS-ImpL2
dilp2>dilp2RNAi
I
J
K
L
P1
DAPI
p<0.0001
p<0.0001
Plasmatocyte differentiation index
100
80
60
40
20
0
wild type
UAS-ImpL2
dilp2RNAi
dilp2-Gal4

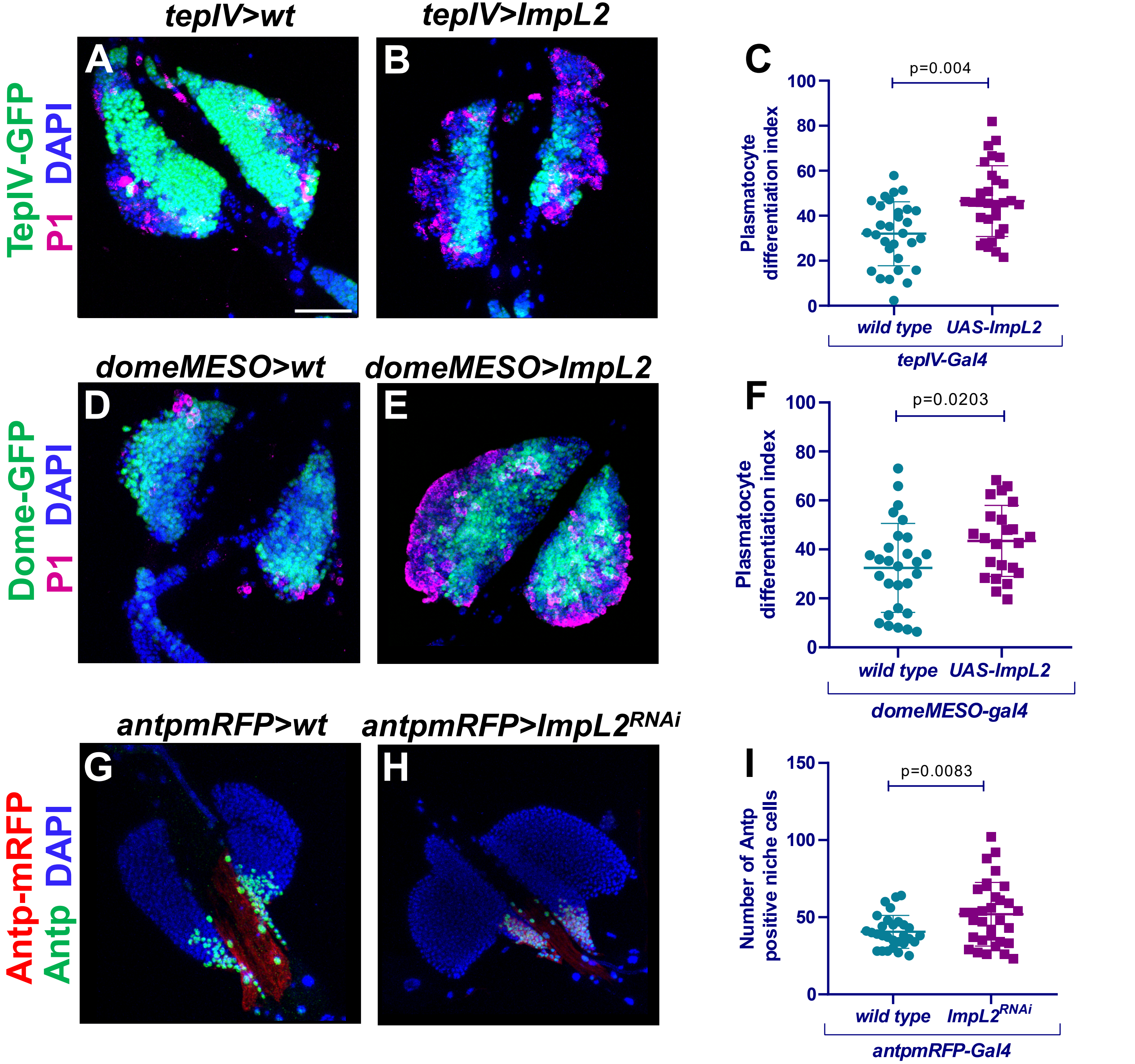

tepIV>wt
tepIV>ImpL2
A
B
C
TepIV-GFP P1 DAPI
p=0.004
Plasmatocyte differentiation index
100
80
60
40
20
0
wild type
UAS-ImpL2
tepIV-Gal4
domeMESO>wt
domeMESO>ImpL2
D
E
F
Dome-GFP P1 DAPI
p=0.0203
Plasmatocyte differentiation index
100
80
60
40
20
0
wild type
UAS-ImpL2
domeMESO-gal4
antpmRFP>wt
antpmRFP>ImpL2RNAi
G
H
I
Antp-mRFP Antp DAPI
p=0.0083
Number of Antp positive niche cells
150
100
50
0
wild type
ImpL2RNAi
antpmRFP-Gal4

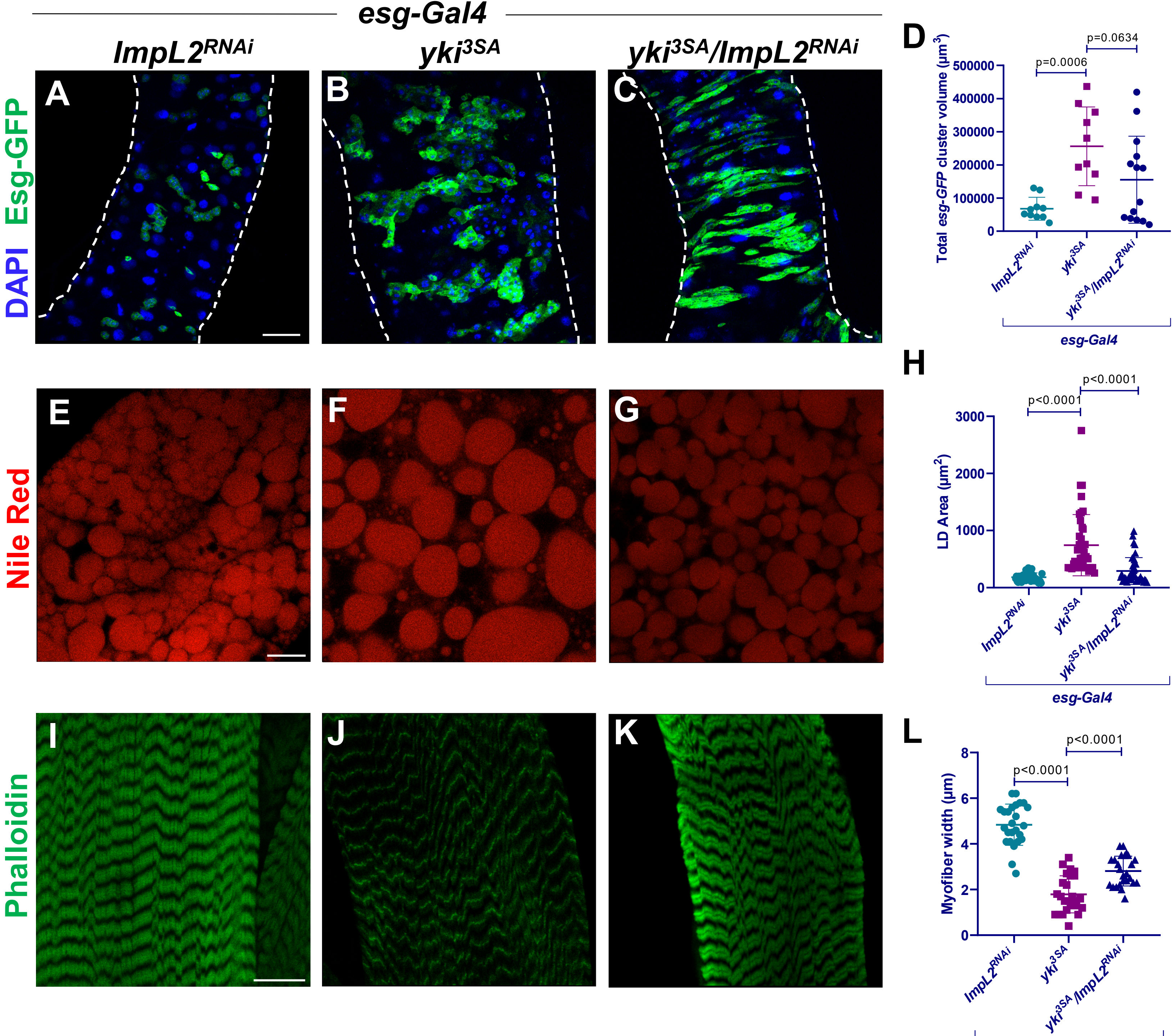
esg-Gal4
ImpL2RNAi
yki3SA
yki3SA/ImpL2RNAi
A
B
C
D
Esg-GFP
DAPI
Total esg-GFP cluster volume (μm3)
p=0.0006
p=0.0634
500000
400000
300000
200000
100000
0
esg-Gal4
E
F
G
H
Nile Red
LD Area (μm2)
p<0.0001
p<0.0001
3000
2000
1000
0
I
J
K
L
Phalloidin
Myofiber width (μm)
p<0.0001
p<0.0001
8
6
4
2
0

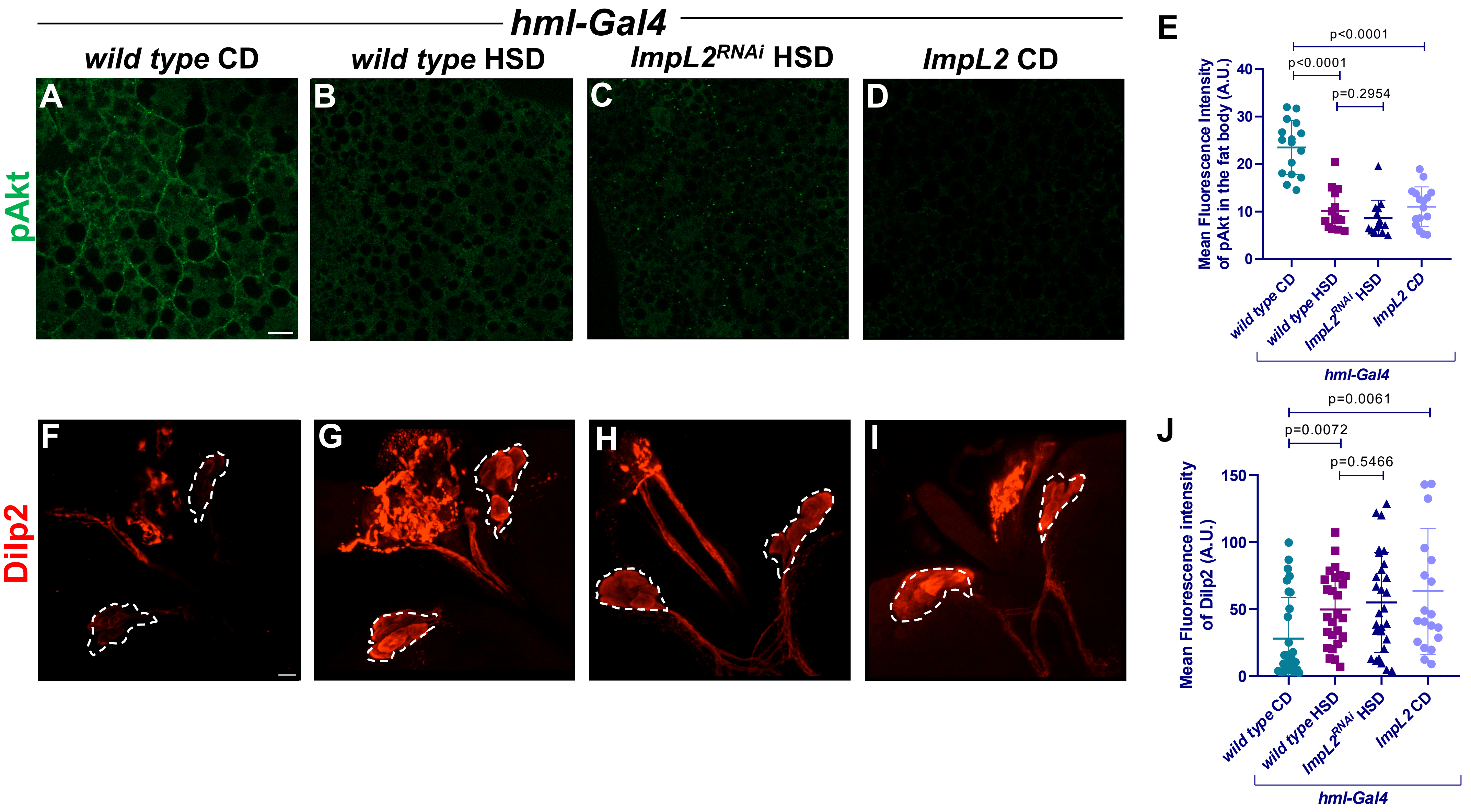

hml-Gal4
wild type CD
wild type HSD
ImpL2RNAi HSD
ImpL2 CD
A
B
C
D
pAkt
E
p<0.0001
p<0.0001
p=0.2954
Mean Fluorescence Intensity of pAkt in the fat body (A.U.)
40
30
20
10
0
wild type CD
wild type HSD
ImpL2RNAi HSD
ImpL2 CD
hml-Gal4
F
G
H
I
Dilp2
J
p=0.0061
p=0.0072
p=0.5466
Mean Fluorescence intensity of Dilp2 (A.U.)
150
100
50
0
wild type CD
wild type HSD
ImpL2RNAi HSD
ImpL2 CD
hml-Gal4

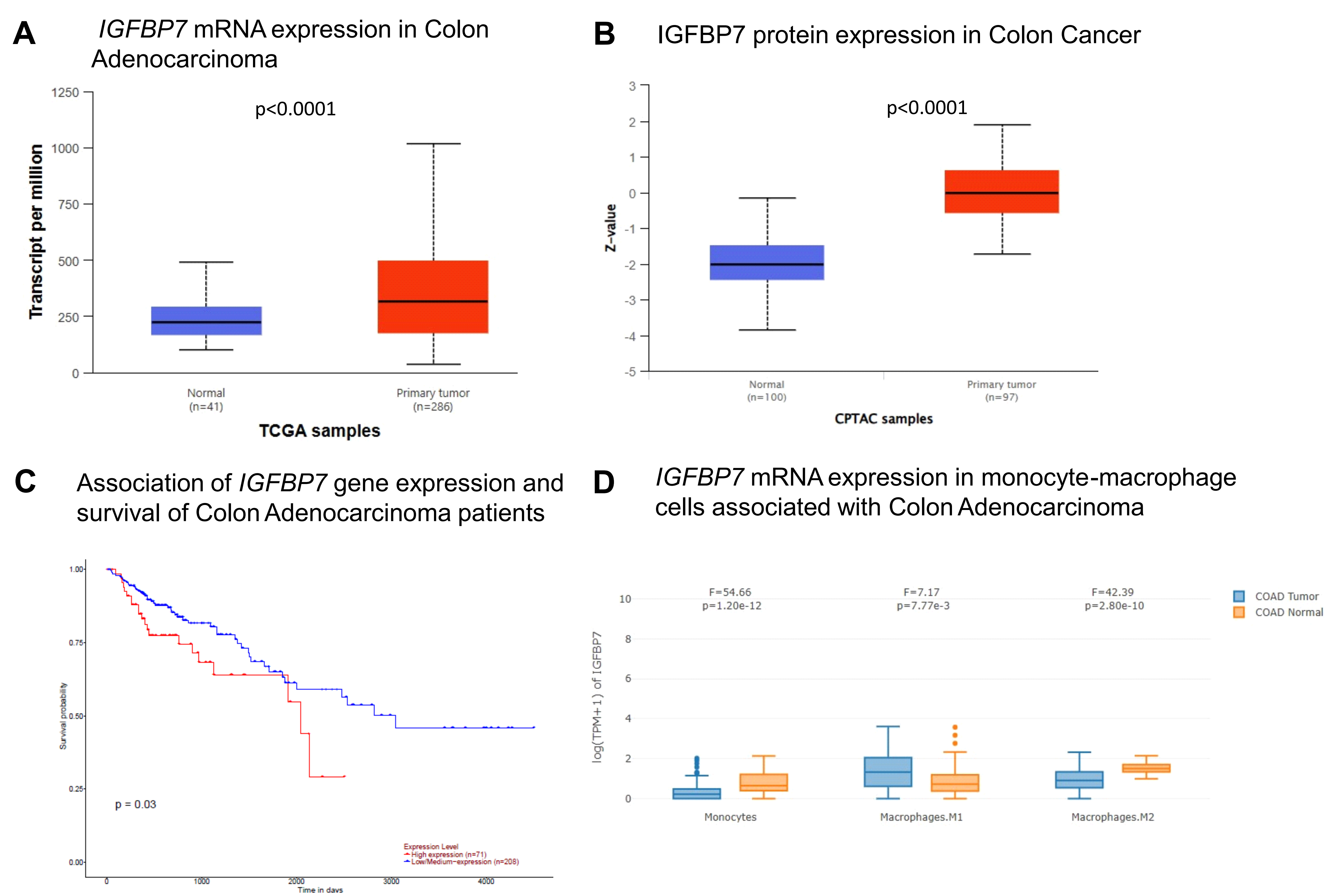
A
IGFBP7 mRNA expression in Colon Adenocarcinoma
p<0.0001
Transcript per million
1250
1000
750
500
250
0
Normal
(n=41)
Primary tumor
(n=286)
TCGA samples
B
IGFBP7 protein expression in Colon Cancer
p<0.0001
Z-value
3
2
1
0
-1
-2
-3
-4
-5
Normal
(n=100)
Primary tumor
(n=97)
CPTAC samples
C
Association of IGFBP7 gene expression and survival of Colon Adenocarcinoma patients
Survival probability
1.00
0.75
0.50
0.25
0.00
p = 0.03
Expression Level
High expression (n=71)
Low/Medium-expression (n=208)
0
1000
2000
3000
4000
Time in days
D
IGFBP7 mRNA expression in monocyte-macrophage cells associated with Colon Adenocarcinoma
log(TPM+1) of IGFBP7
10
8
6
4
2
0
F=54.66
p=1.20e-12
F=7.17
p=7.77e-3
F=42.39
p=2.80e-10
COAD Tumor
COAD Normal
Monocytes
Macrophages.M1
Macrophages.M2
